# Iterative Epigenomic Analyses in the Same Single Cell

**DOI:** 10.1101/2020.07.20.212969

**Authors:** Hidetaka Ohnuki, David J. Venzon, Alexei Lobanov, Giovanna Tosato

## Abstract

Gene expression in individual cells is epigenetically regulated by DNA modifications, histone modifications, transcription factors and other DNA-binding proteins. It has been shown that multiple histone modifications can predict gene expression and reflect future responses of bulk cells to extracellular cues. However, the predictive ability of epigenomic analysis is still limited for mechanistic research at a single cell level. To overcome this limitation, it is useful to acquire reliable signals from multiple epigenetic marks in the same single cell. Here, we propose a new approach for analysis of several components of the epigenome in the same single cell. The new method allows reanalysis of the same single cell. We found that reanalysis of the same single cell is feasible, and provides confirmation of the signals and allows application of statistical analysis to identify reliable signals using data sets generated only from the single cell. Reanalysis of the same single cell is also useful to acquire multiple-epigenetic marks from the same single cells. The method can acquire at least 4 epigenetic marks, H3K27ac, H3K27me3, mediator complex subunit 1 and a DNA modification. We predicted active signaling pathways in K562 single cells using the data. We confirmed that the prediction results showed a strong correlation with actual active signaling pathways shown by RNA-seq results. These results suggest that the new approach provides mechanistic insights for cellular phenotypes through multi-layered epigenome analysis in the same single cells.

## Introduction

A cell can accomplish a number of tasks responding to extracellular and intracellular signals by integrating complex gene-regulatory networks (GRNs) controlled by DNA, the epigenome, RNA and protein (Davidson and Erwin 2006). Emerging single-cell technologies can now measure components of GRNs, including the genome, the transcriptome and the proteome (Efremova and Teichmann 2020). These technologies have opened new and exciting opportunities for deciphering and reconstructing GRNs that drive cell functions (Aibar et al. 2017; Stuart et al. 2019; Stuart and Satija 2019; Welch et al. 2019). Yet, advances in single-cell epigenomic analysis are urgently needed to improve reconstruction of reliable and robust GRNs in single cells.

In bulk cell analysis, characterization of multiple histone modifications predicted gene expression more effectively than characterization of single histone modifications (Karlic et al. 2010; Dong and Weng 2013; Singh et al. 2016; Sekhon et al. 2018; Yin et al. 2019). In addition to capturing concurrent patterns of gene expression, broad epigenomic profiling in bulk cells could predict patterns of gene expression and cell phenotype in response to environmental stimuli (Bock et al. 2011; Krausgruber et al. 2020). These observations suggest that characterization of multiple histone modifications and DNA binding proteins may have a similar potential even at a single-cell level. However, technical limitations in current single-cell epigenomic technologies impede a broad profiling of histone modifications, DNA modifications and DNA binding proteins.

The nucleosome, the basic unit of chromatin structure and epigenetic signaling module, only possesses one double-stranded DNA segment per nucleosome or one binding transcription factor. This limits the number of attainable epigenomic signals per nucleosome resulting in digital/binary-like, low signal patterns. Furthermore, existing single-cell epigenomic technologies cleave genomic DNA and discard the single cells after single use preventing re-analysis of the same single cell to confirm the results and collect data of additional epigenetic marks. To account for these physical limitations, we developed a new single-cell method for epigenomic analysis stemming from the development of a “reusable” single cell. We show that “reusable” single cells afford repeating and extending epigenomic characterization of the same cell. Repeated experiments with the same single-cell revealed that signals from specific antibodies can be reproduced and distinguished from background noise by statistical analysis. Additionally, analysis of different epigenetic markers, such as H3K27ac, H3K27me3, Med1 and 5hmC in the same single cell revealed that it is possible to broaden the epigenomic profile of a single cell. Thus, our new method can capture signals of histone modifications, DNA modification and interacting protein in the same single cell providing an opportunity for integration in GRNs.

## Results

### Method Design: A “Reusable” single cell for EPigenomic analysis (REpi-seq)

The new method consists of two sequential steps. The first step (**Fig. 1A**) generates “reusable” single cells. Cellular proteins, including nuclear proteins, are modified with monomer acrylamide using a paraformaldehyde (PFA)/acrylamide mixture; the monomer acrylamide on the proteins is incorporated into a polyacrylamide scaffold by polymerizing the acrylamide (**Fig. 1A**). Individual cells are then embedded in a polyacrylamide gel-bead (**Supplemental Fig. S1A**). The second step acquires locational information of individual antibodies on the genome through a series of biochemical reactions (**Fig. 1B**). Random primers annealed to the genomic DNA are extended with a DNA polymerase to acquire locational information on the genome. A DNA polymerase, which lacks exonuclease activity, is used to protect genomic DNA. Antibodies are conjugated to a DNA probe containing a unique barcode and a ligation sequence. The antibodies are incubated with the “reusable” single cell. After washing the cell, proximity ligation is performed between the antibody-DNA-probe-conjugate and the random primers using a ligation adapter with a T4 DNA ligase. The ligated products are amplified by MALBAC with a second set of random primers containing a cell-specific barcode. The antibody-DNA-probe-conjugate also contains a T7 promoter sequence. In vitro transcription, RNA purification and reverse transcription are performed to reduce MALBAC byproducts, *i.e.* genome sequences lacking the antibody-DNA probe (**Supplemental Fig. S2**).

**Figure 1.**
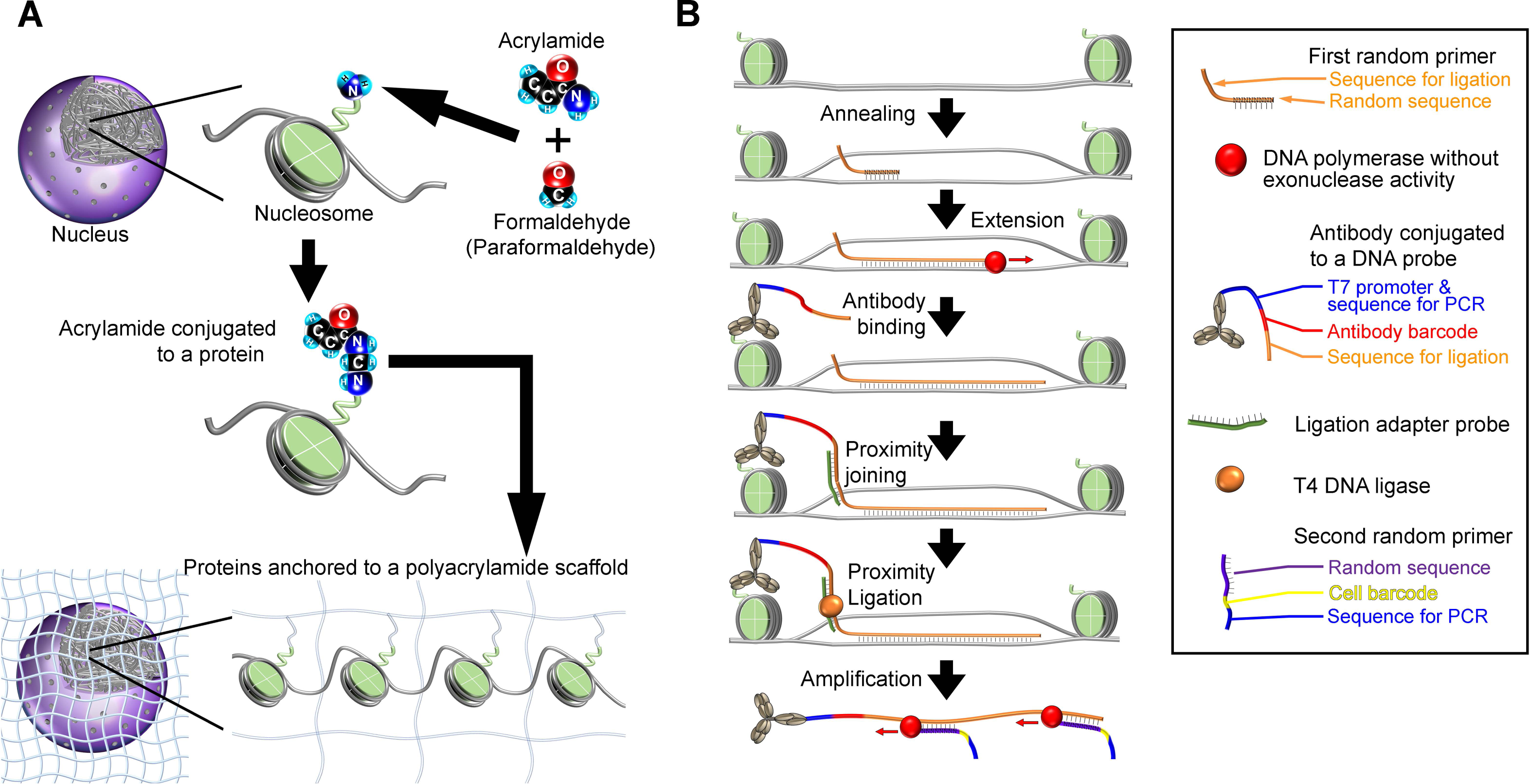
A “Reusable” Single Cell for Epigenomic Analysis (REpi-seq)

### A Single Cell Anchored to a Polyacrylamide Scaffold is Reusable in REpi-seq

We first evaluated whether “reusable” single cells can retain cellular proteins over repeated heating and cooling cycles, which are used for acquiring locational information of antibodies (**Fig. 1B**). The results (**Supplemental Fig. S1B**) indicated that cells treated with 28% acrylamide+4% PFA before embedding into the polyacrylamide scaffold preserve the highest amount of cellular protein over 100 annealing cycles compared to other solutions.

Next, we tested whether REpi-seq can generate the desired products. We performed the series of reactions (**Fig. 1B**) using 8 single cells anchored to the polyacrylamide scaffold, a single cell embedded into the gel without protein anchoring to the gel and blank gel (no cell). Antibodies (anti-H3K27ac and anti-H3K27me3) and control IgG were used in the experiment. Final DNA products are shown in **Supplemental Figure S3A**. Single cells anchored to polyacrylamide generated more DNA products than either blank gel (no cell) or a single cell without anchoring, suggesting that anchoring cellular proteins and genomic DNA to polyacrylamide increases output of desired products. Shallow sequencing using MiSeq (**Supplemental Fig. S3B**) indicated that a single cell anchored to polyacrylamide generates a higher number of desired DNA products containing antibody barcode, ligated sequence and genomic sequences than single cells without anchoring.

### Features of Products Generated by REpi-seq

We analyzed features of the products generated by REpi-seq identified by having sequences of the antibody probe and proximity ligation. Sequencing quality scores of the genomic sequences acquired by the extension of the 1^st^ random primer were above 30 in most reads, indicating less than 0.01% probability of incorrect base calling (**Supplemental Fig. S4**). The length of genomic sequences acquired by the extension of the 1^st^ random was shorter than 32 bp (**Supplementary Fig. 5**). We checked the mapping accuracy of the short genomic sequences (**Supplemental Fig. 6A**). On average, the mapping quality score of the mapped reads was 42.02 and the median was 33. A mapping quality score above 30 indicates that the probability of incorrect mapping is less than 0.1%.In REpi-seq, the ligated products are amplified by MALBAC. We evaluated duplication rates by MALBC in data sets used in the subsequent analyses (**Supplemental Fig. 6B-6D**). MALBAC duplicates were identified using a unique molecular identifier (UMI) of the original ligated products, which comprises the antibody barcode, 8 nt sequence of the 1^st^ random primer and added genomic sequences by the extension. The duplication rates per original ligated products were: 1.106 for anti-H3K27ac derived products, 1.104 for anti-H3K27me3 derived products, 1.187 for anti-Med1 derived products and 1.203 for anti-5hmC derived products. The MALBAC duplicates were removed using the UMI for subsequent analyses.

### REpi-seq has a lower bias for nucleosome-free than for nucleosome-containing regions

REpi-seq acquires locational information of individual antibodies on the genome using the 1^st^ random primer and antibodies in combination with a series of biochemical reactions. We evaluated the presence of bias in REpi-seq products. Nucleosome-free regions have been shown to have higher accessibility by enzymes and molecules(Schones et al. 2008; Gangadharan et al. 2010; Cui and Zhao 2012). We evaluated bias for nucleosome-free regions (**Supplemental Fig 7**). MALBAC showed higher amplification efficiency for longer products of the ligated products than for shorter products (**Supplemental Fig. 7A**). However, we did not observe a positive bias for known nucleosome-free regions in K562 cells (**Supplemental Fig. 7B and 7C**). Rather, the length of nucleosome-free regions was inversely correlated with the number of REpi-seq products.

### Preservation of nuclear proteins position over repeated experiments

Cellular proteins were retained over repeated experiments (**Supplemental Fig. S1B**). Next, we evaluated whether REpi-seq maintains the position of nuclear proteins and genomic DNA over repeated experiments. We repeated the same experiment three times using the same 8 single cells with anti-H3K27ac, anti-H3K27me3 and control IgG. Amplification duplicates were computationally removed based on combinations of antibody barcode, 8 nt random sequence of the 1^st^ random primer and mapped position of the 5’ end on the genome. After removal of amplification duplicates, the read distribution was analyzed in the first, second and third repeated experiment (**Fig. 2A** and **2B**). We identified the signal distribution on the genome detected in the first experiment (green). Subsequently, we identified the signal distribution detected in the second (blue) and the third (red) repeated experiment in the same genomic regions detected in the first experiment. Read distribution patterns in the second and third experiment showed similar patterns to those found in the first experiment. These results indicated that the position of H3K27ac and H3K27me3 marks is closely maintained over the replicate experiments. These results support the conclusion that the polyacrylamide-anchored single cells are reusable at least 3 times.

**Figure 2.**
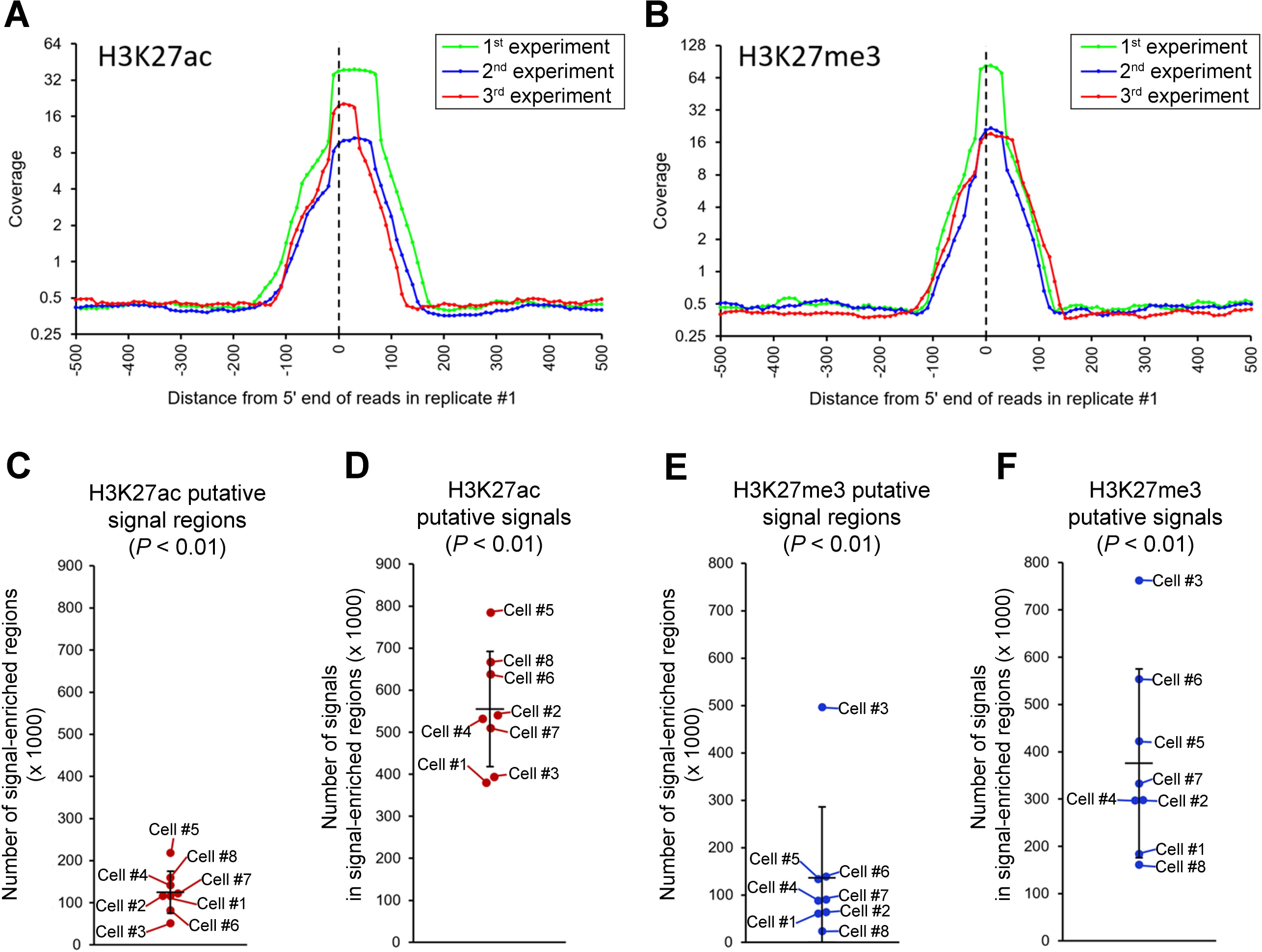
Repeated Experiments Identify Reproduced Signals by Statistical Analysis in the Same Single Cells (A and B) Locations of H3K27ac (A) and H3K27me3 (B) are preserved over repeated experiments in 8 single K562 cells. Genomic regions containing signals in the first experiment were analyzed in the two subsequent experiments. Signal distribution in the first (green line), second (blue line) and third (red line) experiment is shown. (C and E) Number of signal-enriched genomic regions (P < 0.01) in each single cell (no aggregation of data from other single cells). Signal-enriched regions were identified by bootstrap statistical test based on the different distribution of antibody signals from randomized antibody signals and control lgG (see Supplemental Methods for details). (D and F) Number of signals per cell from antibodies to H3K27ac (D) and H3K27me3 (F). Putative signals (P < 0.01) were counted in all signal-enriched genomic regions in each cell. Horizontal bar: mean of 8 single cells. Error bars: standard deviation.

### Repeated Experiments Identify Reproduced Signals by Statistical Analysis in the Same Single Cells

We further analyzed data from the 3 repeated experiments using the same reusable single cells to answer the question of whether signals from specific epigenetic antibodies differ from those of control IgG and stochastic background. To identify genomic regions where antibody-derived signals are enriched compared to control IgG and stochastic noise, we applied the bootstrap statistical test (see supplemental Methods) to each 500 bp genomic region (shifted every 250 bp) from each of the triplicate experiments with each single cell. The test results identified, on average, 125,502 signal-enriched regions containing H3K27ac signals (**Fig. 2C**) and 136,387 signal-enriched regions containing H3K27me3 signals (**Fig. 2E**). These regions were statistically significantly different from controls (p < 0.01). These signal-enriched regions contained, on average, 556,432 H3K27ac signals per cell (**Fig. 2D**) and 375,756 H3K27me3 signals per cell (**Fig. 2F**). Single-cell #3 was an outlier for H3K27me3 (**Fig. 2E** and **2F**). However, single-cell #3 showed an inverse trend for H3K27ac (**Fig. 2C** and **2E**), suggesting that results from single-cell #3 are not reflective of experimental error. Signals in the enriched 500 bp regions amounted, on average, to 4.91 H3K27ac signals and 3.81 H3K27me3 signals. Since a nucleosome includes 146 bp of wrapping DNA and the length of linker DNA is up to 80 bp, a 500 bp genome region may contain 2 to 3 nucleosomes. Since a nucleosome contains 2 copies of histone H3, 4 to 6 copies of histone H3 may be present per 500 bp of the genome. Therefore, the average frequencies of 4.91 H3K27ac and 3.81 H3K27me3 signals per 500 bp region, detected here likely reflect high-density signal regions. These results also indicate that repeated experiments using the same single cells accumulate signals in specific regions compared to controls. In a single experiment, one nucleosome of a cell can generate only a few signals because one nucleosome has only one dsDNA. In repeated experiments, the results revealed signal patters from epigenetic antibodies that are distinct from the random pattern of controls. Statistically significant (p < 0.01 by the bootstrap test) signals were considered putative signals in the subsequent analyses.

### Evaluation of Putative Signals at Promoters and Enhancers

Since histone H3K27ac is enriched at promoter and enhancer regions (Creyghton et al. 2010), we evaluated the location of putative single-cell H3K27ac signals in promoters and enhancers by comparing with bulk ChIP-seq results (**Fig. 3A-3D**). For this analysis, we used the Human ACtive Enhancers to interpret Regulatory variants (HACER) atlas (Wang et al. 2019), which defines cell-specific, active enhancers based on bidirectional enhancer RNA expression in 265 human cell types. HACER also contains experimentally validated data based on active enhancers-promoter interaction. In HACER, enhancers and promoters are classified into 2 groups, K562-cell-type specific, active enhancer/promoter (referred here as K562-active enhancer/promoter) and K562-cell-type non-specific, active enhancer/promoter (referred here as other active enhancer/promoter). REpi-seq detected 9,974 promoters with putative signal enrichment (*p* < 0.05), 62.93% of which were confirmed by bulk ChIP-seq (**Fig. 3A**). Among the K562-active promoter (**Fig. 3B**), REpi-seq detected 4,212 promoters, 77.28% of which were confirmed by bulk ChIP-seq. Among other active enhancers (**Fig. 3C**), REpi-seq detected 78,874 enhancers with signal enrichment (p < 0.05), 25.77% of which were confirmed by bulk ChIP-seq. Among K562-active enhancers detected by REpi-seq (**Fig. 3D**), 73.89% were confirmed by bulk ChIP-seq. The 73.89% confirmation rate of K562-active enhancers is higher than the 25.77% confirmation rate of other active enhancers. These results indicate that putative signals identified by REpi-seq and the bootstrap statistical test detect established cell-specific epigenetic signals. These results further suggest that REpi-seq can recognize other, cell-type non-specific, active enhancers in K562 cells.

**Figure 3.**
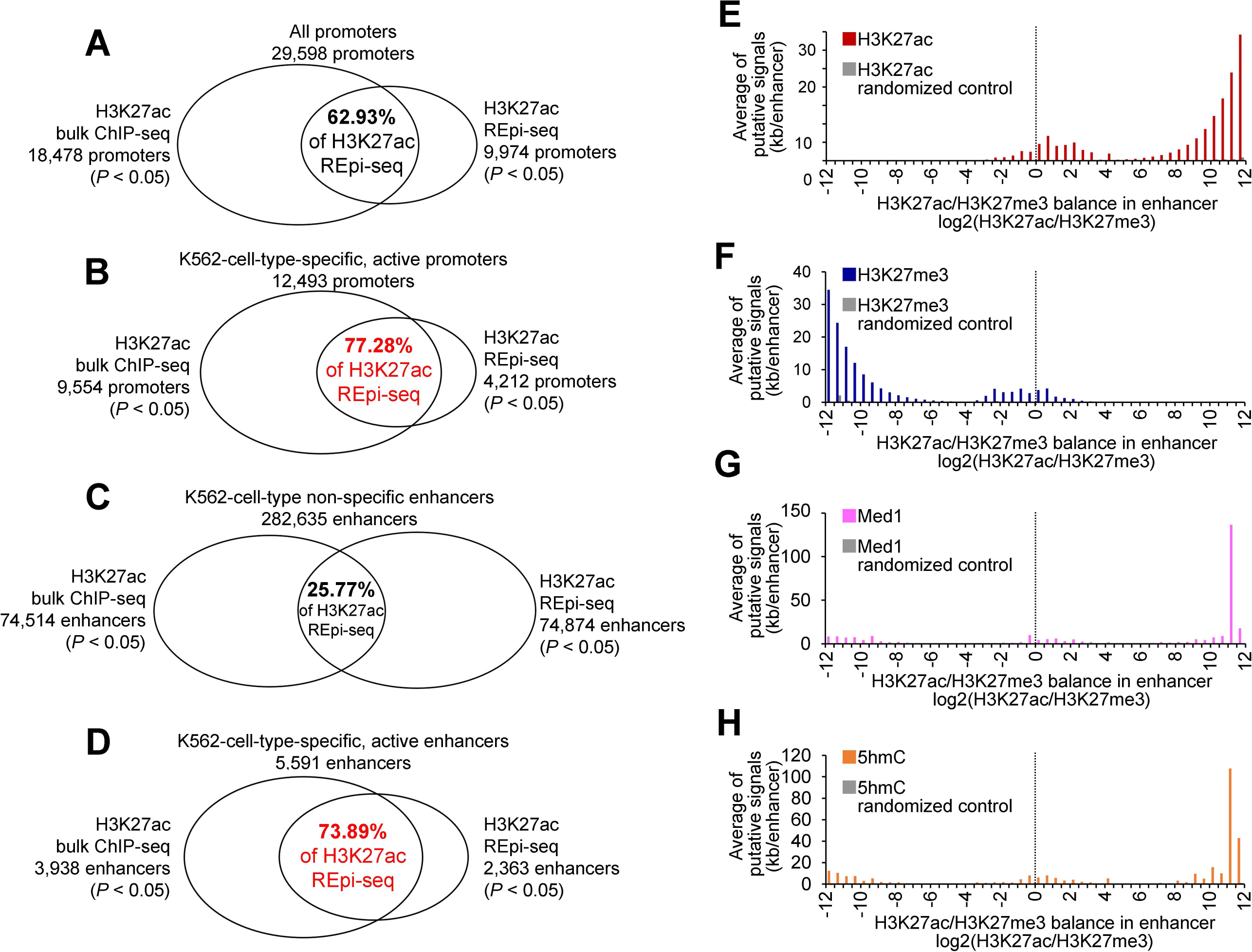
Validation of REpi-seq Putative Signals (A-0) Validation of REpi-seq putative signals by bulk ChlP-seq. H3K27ac signal enrichment in all promoters (A), K562-cell-type-specific, active promoters (B), K562-cell-type non-specific, active enhancers (C), K562-cell-type-specific, active enhancers (D).Statistical significance of signal enrichment in promoters and enhancers was calculated by bootstrap statistical test. Signal enriched promoters and enhancers (P < 0.05) are shown in the Venn diagrams (see STAR Methods for details). (E-H) Validation of REpi-seq putative signals by enhancer classification. Enhancers were classified by the relative ratio Log2(H3K27ac/H3K27me3,) as detailed in Online Methods. H3K27ac (E), H3K27me3 (F), mediator complex subunit 1 (Med1, G) and 5-hydroxymethylcytosine (5hmC, H) signal counts in the classified enhancers are shown in the Y-axis.

### Enhancer Classification using Four Epigenetic Marks

The presence of H3K27ac uniquely marks active enhancers that are linked to active genes (Creyghton et al. 2010; Rada-Iglesias et al. 2011). Poised or inactive enhancers that are linked to inactive genes are distinguished by the absence of H3K27ac and enrichment of H3K27me3 (Rada-Iglesias et al. 2011). Ratios of H3K27ac/H3K27me3, but not simply by presence or absence of H3K27ac or H3K27me3, explain changes in gene expression during hematopoietic lineage differentiation (Prescott et al. 2015; Gray et al. 2017). Here, we used relative ratios Log2(H3K27ac/H3K27me3) (shown in the X-axis, **Fig. 3E-3H**) to classify the enhancers. Results are expressed as average number of H3K27ac (Y-axis, **Fig. 3E**) and H3K27me3 (Y-axis, **Fig. 3F**) putative signals per kb/enhancer. Consistent with previous reports (Creyghton et al. 2010; Rada-Iglesias et al. 2011), enhancers with Log2(H3K27ac/H3K27me3) values above 6 were exclusively H3K27ac-positive and H3K27me3-negative. In the transitional −5 to +5 range, H3K27ac signals were dominant in the 0 to +5 range and H3K27me3 signals were dominant in the 0 to −5 range. In the less than −5 range, H3K27me3 was exclusively positive in enhancers. These results indicated that signals from REpi-seq could classify enhancers into at least 3 groups; poised enhancers (exclusively H3K27me3-positive), intermediate enhancers (transitional) and active enhancers (exclusively H3K27ac-positive).

Super-enhancers are clusters of highly active enhancers marked by high-level H3K27ac, high-density master transcription factors and mediator complexes (Whyte et al. 2013). Therefore, if the enhancer classification by REpi-seq (**Fig. 3E** and **3F**) is correct, mediator complex subunit1 (Med1) should be enriched in the active enhancers. To examine this, we re-tested the same 8 single cells with anti-Med1 antibody and control IgG (4^th^ experiment). Med1 putative signals were identified by bootstrap test statistics using aggregated datasets from 8 single cells (p < 0.01, **Supplemental Fig. S8A** and **S8B**). This aggregated data identified putative common Med1 signals among the 8 single cells based on signal-enriched regions over control IgG and stochastic noise. The results from counting Med1 signals in the enhancers classified by REpi-seq (**Fig. 3G**) show that Med1 signals are enriched in the active enhancers, especially in highly active enhancers. These results also indicate that REpi-seq data can correctly identify the epigenetic status of enhancers as defined by the relative ratio of Log2(H3K27ac/H3K27me3).

We evaluated further the enhancer classification using the additional epigenetic mark, 5-hydroxymethylcytosine (5hmC). Stroud et al. (Stroud et al. 2011) reported that 5hmC is significantly enriched in enhancers with H3K27ac. In addition, 5hmC is generated by the enzyme Ten-eleven translocation 1 (TET1) (Ito et al. 2011; Williams et al. 2011) that interacts with Med1 super-enhancer complex (Ding et al. 2015). If the enhancer classification by REpi-seq was correct, 5hmC should be enriched in active enhancers. For this evaluation, we re-tested the same 8 single cells with anti-5hmC and control IgG (5^th^ experiment). Putative signals of 5hmC were identified based on statistical significance (bootstrap test; p < 0.01) using aggregated datasets from 8 single cells (**Supplemental Fig. S8C** and **S8D**). Next, we calculated the average number of putative 5hmC signals in each classified enhancer range (**Fig. 3H**). Consistent with previous reports(Stroud et al. 2011), the results indicate that the 5hmC putative signals are enriched in the active enhancers. These results also indicate that the H3K27ac and H3K27me3 putative signals from REpi-seq can correctly recognize the epigenetic status of enhancers. Furthermore, these results also indicate that the reusable single cells are reusable at least 5 times in REpi-seq, and that REpi-seq can analyze at least 4 epigenetic marks in the same single cells.

### Signal Evaluation in Enhancers Based on Gene Targets

The results of REpi-seq testing provide evidence that H3K27ac and H3K27me3, combined, can correctly identify active enhancers in K562 single cells. We now assessed the function of these active enhancers by focusing on interacting genes. The HACER dataset (Wang et al. 2019) contains experimentally validated interactions between enhancers and genes from 4DGenome (Teng et al. 2015) and chromatin interaction studies from Hi-C, ChIA-PET, HiChIP and Capture HiC. K562-active enhancers detected by REpi-seq (2,363 enhancers) and by bulk ChIP-seq (3,938 enhancers) interact with 3,154 and 5,002 genes, respectively (**Fig. 4A**). Interacting genes identified by REpi-seq were mostly (93.18%) common with those identified by bulk ChIP-seq. Among other active enhancers (**Fig. 4B**), overlap between REpi-seq and bulk ChIP-seq (78.44%) was smaller than with the K562-active enhancers (93.18%). Since enhancer-promoter/gene interactions transmit enhancer activity to target genes (Robson et al. 2019), these results suggest that signals from REpi-seq can correctly capture sets of enhancers that regulate functions of K562 single cells.

**Figure 4.**
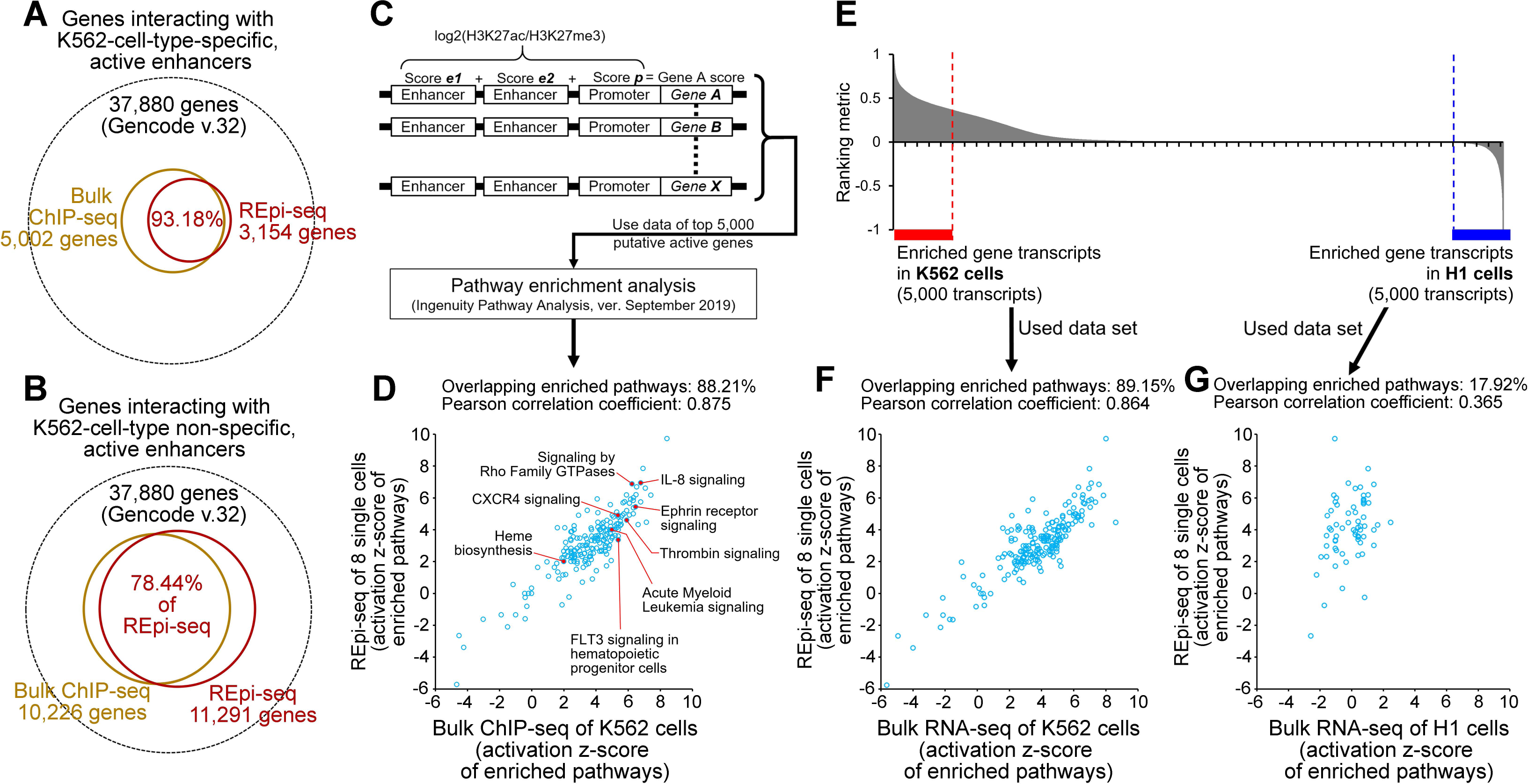
Evaluation of REpi-seq Putative Signals by Target Genes and Target Pathways (A and B) Genes interacting with K562-cell-type-specific, active enhancers (A) and K562-cell-type non-specific, active enhancers (B) in K562 cells, are shown. The activity of enhancers was determined by the relative ratio Log2(H3K27ac/H3K27me3) from bulk ChlP-seq and REpi-seq data, as detailed in the STAR Methods. (C) Process for transmission of promoter+enhancers activity to a gene and extrapolating the biological function of active enhancers and promoters. (D) Enriched canonical pathways identified by bulk ChlP-seq and REpi-seq. Activation z-scores from ChlP-seq (x-axis) and REpi-seq (y-axis) are shown; Pearson correlation coefficient was calculated. (E) Differential gene expression in bulk K562 and H1 cell by RNA-seq. The top 5,000 transcripts in K562 and H1 cells were used as input data in F and G. (F and G) Enriched canonical pathways identified by REpi-seq (8 single K562 cells) and RNA-seq from bulk K562 (top 5,000 transcripts, F) and by REpi-seq (8 single K562 cells) and RNA-seq from bulk H1 ChlP-seq (top 5,000 transcripts, G).

### Signal Evaluation in Enhancers and Promoters Based on Canonical Signaling Pathways

We further evaluated signals from REpi-seq by pathway enrichment analysis (Kramer et al. 2014) by focusing on analysis of genes proximal to active enhancers and promoters detected by REpi-seq and bulk ChIP-seq. Since enhancer-promoter/gene interactions transmit enhancer activity to target genes(Robson et al. 2019), we mimicked this transmission in silico. We calculated the combined relative scores {Log2(H3K27ac/H3K27me3)} of enhancers and promoters and transmitted this score to the proximal gene (**Fig. 4C**). The transmitted scores to genes were used in pathway enrichment analysis. By comparing the 5,000 genes with the highest scores (putatively active genes) from REpi-seq and bulk ChIP-seq, we found that 88.21% of the enriched canonical pathways (referred here as pathways) from REpi-seq were common to the enriched pathways from bulk ChIP-seq (**Fig. 4D**). Activation *z*-scores of the enriched pathways from REpi-seq and bulk ChIP-seq showed a strong correlation (Pearson correlation coefficient, *r* = 0.875). We further evaluated the results of pathway enrichment in K562 cells based on REpi-seq and RNA-seq of bulk cells. The RNA-seq data from the embryonic stem cell line H1 were used as a reference control. The top 5,000 transcripts from bulk K562 cells and bulk H1 cells were used in pathway enrichment analysis (**Fig. 4E**). Most (89.15%) of the enriched pathways from REpi-seq were confirmed as enriched pathways from bulk RNA-seq of K562 cells (**Fig. 4F**). Activation *z*-scores of the enriched pathways from RNA-seq and REpi-seq showed a strong correlation (Pearson correlation coefficient, *r* = 0.864). However, only 17.92% of the enriched pathways from REpi-seq overlapped with the enriched pathways of bulk H1 RNA-seq, and the overlapped pathways were poorly correlated (Pearson correlation coefficient, *r* = 0.365). These results suggest that signals from REpi-seq are useful to infer downstream functions of epigenetically active enhancers.

### Epigenetic Signature Generated by REpi-seq

The results of Figure 4 supported the conclusion that signals from REpi-seq reflect activation at cis-elements, gene targets of cis-elements and pathways linked to putative active genes. Next, we evaluated at a cellular level the signals generated by REpi-seq. To this end, we utilized the epigenetic status [Log2(H3K27ac/H3K27me3)] of enhancers and promoters to compare cell types stochastic neighbor embedding (t-SNE, **Fig. 5A**). K562 single cells (red dots) and K562 bulk cells (blue dots) differed from endothelial cells (yellow), fibroblasts (green) and myoblasts (light blue). The K562 single and bulk cells bordered hematopoietic malignant cell lines and primary hematopoietic cells. K562 cell line is a myelogenous leukemia cell line (Lozzio and Lozzio 1975) with multi-differentiation potential along erythroid, macrophage and megakaryocytic lineages (Chylicki et al. 2000; Duncan et al. 2016; Yang et al. 2016). Contour plots of K562 single cells (red lines) and K562 bulk cells (blue lines) indicated epigenetic similarities between datasets from K562 single and bulk cells. These data support the conclusion that signals generated by REpi-seq can capture cell-type specific epigenetic signatures at a cellular level.

**Figure 5.**
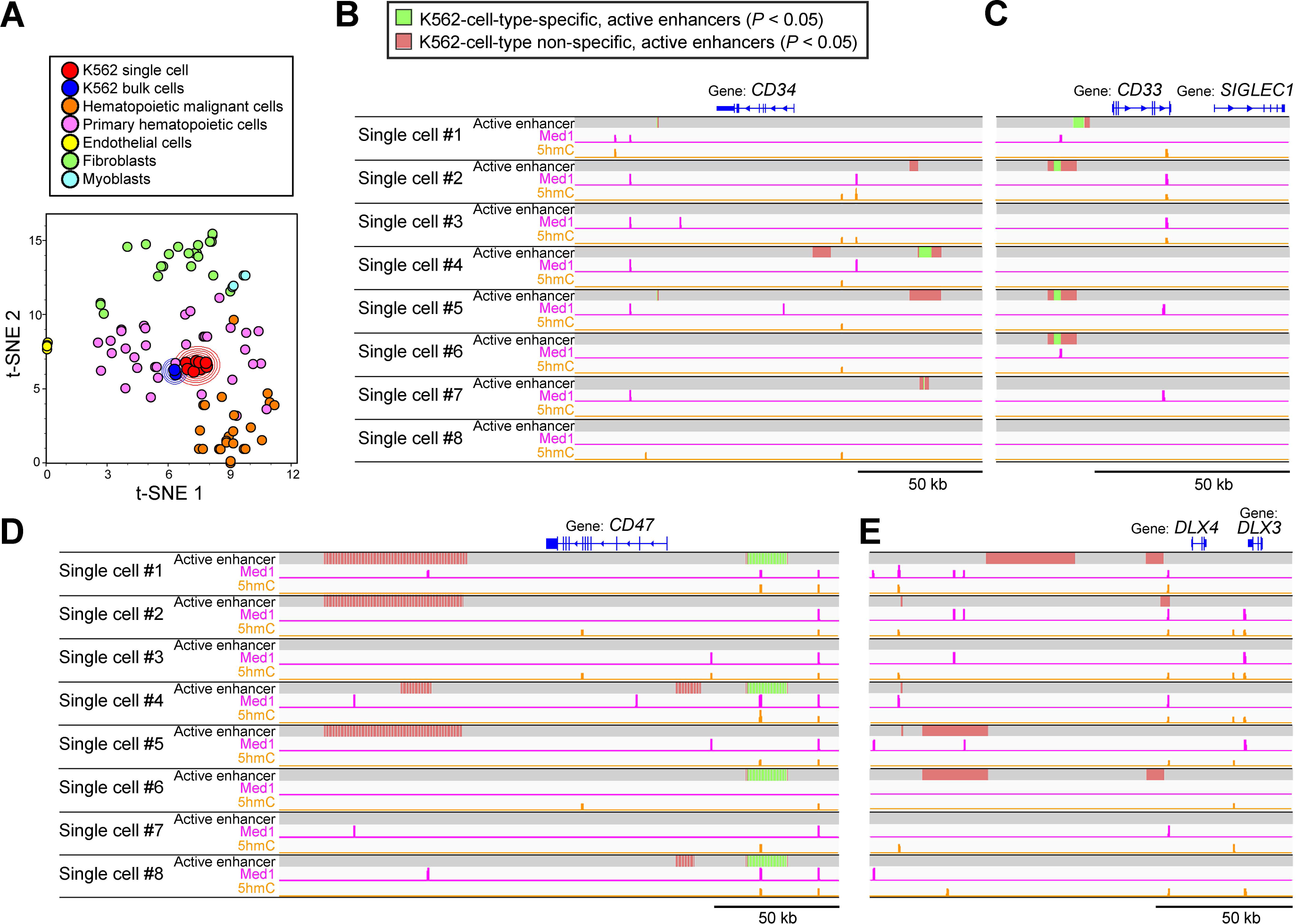
Visualization of Epigenetic Signatures Generated by REpi-seq using Dimensionality Reduction 50 kb (A) Dimensionality reduction of whole data sets generated by REpi-seq and bulk ChlP-seq. Epigenetic status of enhancers and promoters was determined from bulk ChIP-seq data of various cell types and from REpi-seq of 8 K562 cells by Log2(H3K27ac/H3K27me3), as detailed in the Online Methods. Dimensions of cell data were compressed by t-Stochastic Neighbor Embedding (t-SNE). (B-F) Identified active enhancers, Med1 and 5hmC signals by statistical analysis. Statistically significant activation of enhancers was determined by bootstrap test (P < 0.05) using data sets of H3K27ac and H3K27me3 generated by REpi-seq. Identified active enhancers were further classified into K562-cell-type-specific, active enhancers (green) and K562-cell-type non-specific, active enhancers (red) by bulk ChIP-seq data. Med1 and 5hmC signals (P < 0.01) were also identified by statistical analysis, bootstrap test. Loci of CD34 (B), CD33 (C), CD47 (D) and DLX4 (E) are shown. See also Figure S5.

### Active Enhancers Usage and Med1 Association among Single Cells

**Figure 3C-3H** shows the association of Med1 and 5hmC with active enhancers. K562 cells have features shared by common myeloid progenitors (CMPs), megakaryocyte-erythroid progenitor (MEPs) and/or granulocyte-monocyte progenitor (GMPs), and display differentiation potential along the erythroid, myeloid and megakaryocytic lineages (Klein et al. 1976; Andersson et al. 1979), (Lozzio et al. 1981). CD34 is expressed in CMP and GMP, but not in MEP (Warren et al. 2006). Stem-like chronic myeloid leukemia cells are marked by CD33 and CD47 (Valent et al. 2014). The epigenetic status of the CD34 locus, the CD33 locus and the CD47 locus in the 8 K562 single cells is shown in **Figure 5B-5D**. K562-active enhancers (green) and other active enhancers (red) were detected in some of the single K562 cells by REpi-seq. In addition, Med1 and 5hmC marked active enhancers and other regions in most cells, suggesting the occurrence of interaction between some of the active enhancers and mediator complex. These results indicate that the *CD34*, *CD33* and *CD47* genes could be epigenetically active in some of K562 cells.

Megakaryocytic differentiation is induced in K562 cells by the homeobox transcription factor DLX4 whereas erythroid differentiation is repressed. Other active enhancers were detected upstream of the DLX4 locus, some of which were marked with Med1 (**Fig. 5E**), suggesting an epigenetic bias toward megakaryocytic differentiation in some of the K562 cells.

K562 cells have been used as a Human Leukocyte Antigen (HLA)-deficient cell line to measure cytotoxicity of natural killer cells (Nagel et al. 1981). Expression of *HLA* genes can be induced in K562 cells by sodium butyrate, which inhibits histone deacetylation and increases histone acetylation (Sutherland et al. 1985). Our analysis of *HLA-A*, *HLA-B* and *HLA-C* loci (**Supplemental Fig. S9A-S9C**) is consistent with previous reports (Nagel et al. 1981; Sutherland et al. 1985) in showing that these regions are mostly epigenetically inactive, except for upstream regions of HLA-B in single-cells #5 and #6. Overall, the results in **Figure 5B-5E** and **Supplemental Figure S9A-S9C** indicate that REpi-seq can identify the activation status of cis-elements, association of DNA binding protein, and DNA modification in the same single cell.

## Discussion

Here, we provide initial evidence that the reusable single cell is amenable to re-analysis at least 5 times. Reusable single cells can preserve nuclear proteins and genomic DNA location over repeated experiments. REpi-seq is a useful single-cell method for identification of epigenetic marks at the level of cis-elements (Fig. 3), cis-elements target genes, gene-induced downstream pathways (Fig. 4), and at a cellular level (Fig. 5A).

This method is in its infancy and has limitations. The experiments presented here were performed manually, without automation, as a proof-of-principle. Automation using robots, microfluidics or other innovative approaches is needed to scale up the throughput. Despite such limitation, the results of our experiments suggest the importance of the new method. Thus far, it has been challenging to increase the number of histone modifications and DNA binding proteins analyzed in the same single cell. Our results indicate that at least 4 epigenetic marks could be analyzed in the same single cell. In bulk cells, multiple histone modifications have been shown to improve prediction of gene expression using deep learning approaches (Karlic et al. 2010; Dong and Weng 2013; Singh et al. 2016; Sekhon et al. 2018; Yin et al. 2019). Also, epigenetic profiles can predict bulk cell responses to extracellular cues (Bock et al. 2011; Krausgruber et al. 2020). Broad epigenetic analyses can also provide mechanistic insights into cellular phenotypes. Currently, analysis of epigenetic mechanisms of gene expression and cell responses is challenging in single cells. The current results suggest that analysis of multiple epigenetic marks is possible at a single cell level and may allow predicting gene expression and cellular responses in single cells.

## Methods

Methods are described in supplemental Methods.

## Acknowledgments

We thank Drs. David Sanchez-Martin and Christopher B. Buck for comments during the conceptualization stage of the project. We also thank the Genomics Core, Center for Cancer Research, National Cancer Institute, National Institutes of Health for help in preliminary experiments, and the Collaborative Bioinformatics Resource, CCR, NCI, NIH for advice in computational analysis. We thank Ms. Anna Word for helping with the optimization of DNA polymerases used in the method. This work utilized the computational resources of the NIH HPC Biowulf cluster (http://hpc.nih.gov). This project is supported by the Intramural Program of the Center for Cancer Research, National Cancer Institute, National Institutes of Health, the NCI Director’s Innovation Award (#397172) and Federal funds from the National Cancer Institute under Contract No. HHSN261200800001E. We thank Drs. Tom Misteli, Carol Thiele, Douglas R. Lowy and all members of Laboratory of Cellular Oncology for productive comments. We also thank Drs. Christopher B. Buck, Robert Yarchoan, and Douglas R. Lowy for reviewing the manuscript.

## Author Contributions

Conceptualization, H.O.; Methodology, H.O., Codes, H.O. and A.L.; Validation, H.O. and G.T.; Formal Analysis, H.O. and D.J.V.; Investigation, H.O.; Data Curation, H.O. and A.L.; Writing – Original Draft, H.O.; Writing – Review & Editing, H.O., D.J.V. A.L. and G.T.; Supervision, H.O. and G.T.; Project Administration, H.O. and G.T.; Funding Acquisition, H.O. and G.T.

## Data and Code Availability Data

All sequencing data is available from the NIH National Center for Biotechnology Information Sequence Read Archive (PRJNA602948) at https://dataview.ncbi.nlm.nih.gov/object/PRJNA602948?reviewer=bnn2d3vci3dvf295p985596dlg

The data is also available from Mendeley at https://data.mendeley.com/datasets/zfyxxs382m/draft?a=bb5dbb3c-ee31-4d1d-ab57-6e4cb42be474

## Code

All codes used here are available at https://github.com/HidetakaOhnuki-NCI/REpi-seq. Workflow and data flow of the Shell scripts are shown in the PDF file available at https://github.com/HidetakaOhnuki-NCI/REpi-seq/blob/master/000_Workflow_page1-11.pdf.

## Supplemental material

### Supplemental Methods

#### Cell Culture

K562 cells from the American Type Culture Collection (ATCC, CCL-243) were cultured in Iscove’s Modified Dulbecco’s Medium (12440053, Thermo Fisher Scientific) containing 10% fetal bovine serum (F2442-500ML, Sigma-Aldrich). Cells tested mycoplasma-negative by a PCR-based method (Young et al. 2010).

#### Antibody Conjugation to DNA Probe

Anti-H3K27ac (RRID:AB_2561016, ENCODE ID: ENCAB000ADS, 39133, Active Motif), anti-H3K27me3 (RRID:AB_2561020, ENCODE ID: ENCAB000ADT, 39155, Active Motif), anti-Med1 (RRID:AB_946345, ab60950, Abcam), anti-5hmC (A-1018, Epigentek) antibodies and control IgG (I5006, Sigma-Aldrich) were conjugated with the DNA probes (Integrated DNA technologies) listed in **Table S1**. Glycerol and sodium azide were removed from all antibodies using Zeba Spin Desalting Columns according to the manufacturer’s protocol (89682, Thermo Fisher Scientific).

The antibodies were activated by introducing the heterobifunctional crosslinker Succinimidyl-6-hydrazino-nicotinamide (S-HyNic, S-1002-105, TriLink Biotechnologies) according to the following procedures: S-HyNic (1 mg) was dissolved with 100 μl of anhydrous Dimethylformamide (DMF, S-4001-005, TriLink Biotechnologies), and 0.6 μl of S-HyNic/DMF was added to 100 μl of antibody solution (1 mg/ml). The antibody was incubated for 2 hours at room temperature. Extra S-HyNic was removed using the Zeba Spin Desalting Column.

To conjugate the DNA probe to the activated antibody, the heterobifunctional crosslinker N-succinimidyl-4-formyl benzamide (S-4FB, S-1004-105, TriLink Biotechnologies) was introduced into the DNA probe according to the procedure below. The DNA probes listed in **Table S1** were synthesized with an amino group at 5’ end with a C12 spacer (Integrated DNA Technologies). Twenty nanomoles of the DNA probe was dissolved into 20 μl of 100 mM sodium phosphate/150 mM NaCl, pH8.0. S-4FB (1 mg) was dissolved into 50 μl of anhydrous DMF. Ten μl of S-4FB/DMF was added to the DNA probe and incubated for 2 hours at room temperature. Extra S-4FB was removed using the Zeba Spin Desalting Column.

The activated antibody and the DNA probe were mixed and incubated overnight at room temperature. Unbound DNA probe was removed by Ultrafiltration using Amicon Ultra (Molecular Cut Off 100 kDa, UFC5010, Millipore), and the buffer was exchanged to TBS containing 50% glycerol, 5 mM EDTA and 5% BSA. Antibody concentration is measured by Sandwich ELISA (Weiland and zu Dohna 1978) using a standard curve of rabbit IgG, and antibody concentration was adjusted to 1 mg/ml. The antibody conjugated to the DNA probe was stored at −20°C until use.

#### Preparation of “Reusable” Single Cells

K562 cells were harvested by centrifuging at 1,200 x g for 5 min and washed with PBS. The cells were treated with one of the following solutions: PBS only, PBS containing 4% paraformaldehyde (PFA, 15713, Electron Microscopy Sciences), PBS containing 5% acrylamide and 4% PFA, PBS containing 20% acrylamide and 4% PFA, or PBS containing 28% acrylamide and 4% PFA. After 1-hour incubation, the cells were washed with TBS containing 2.5% BSA. Individual cells were transferred into a 0.2 ml tube containing 3 μl/tube of the acrylamide solution [PBS containing 3.88% (w/v) Acrylamide (01697-500ML, Sigma-Aldrich), 0.12% (w/v) Bis-acrylamide (M1533-25ML, Sigma-Aldrich) and 1% (w/v) ammonium persulfate (A3678-25G, Sigma-Aldrich)] using ALS CellCelector (ALS Automated Lab Solutions) or PicoPipet with a micromanipulator (Nepa Gene) under microscopy. Fifty l of mineral oil containing 0.2% TEMED was added to the 0.2 ml tube. Polymerization was performed at room temperature for 1 hour. A cell was embedded into the outside layer of the two-layered acrylamide bead. The cell was stained with SYBR Gold (S11494, Thermo Fisher Scientific), and the presence of a single cell was confirmed using fluorescent microscopy (BZ-X710, Keyence).

#### Cellular Protein Retention Assay

K562 cells were harvested and washed in PBS. Cells were suspended in PBS and 0.33 x10^5^ cells were aliquoted into 0.5 ml Protein LoBind tubes (0030108094, Eppendorf). The tubes were centrifuged at 1,200 x g for 5 min. As input controls, 1×10^5^ cells were lysed with 100 μl/tube of 1 x lithium dodecyl sulfate buffer (LDS buffer, NP0007, Thermo Fisher) and stored at 4 °C until measurement. The cells were treated with PBS only, PBS containing 4% PFA, PBS containing 5% acrylamide and 4% PFA, PBS containing 20% acrylamide and 4% PFA, or PBS containing 28% acrylamide and 4% PFA. After 1-hour incubation, cells were washed with TBS containing 10% fetal bovine serum. Cells were suspended with 90 μl of the acrylamide solution and the cell suspension was transferred into 9 PCR tubes containing 50 μl/tube mineral oil containing 0.2% TEMED. After 1-hour incubation, polymerized beads (3 beads, approximately 1 x 10^5^ cells) were transferred into a PCR tube. ThermoPol Mg (-) buffer (100 μl/tube) was added and the following treatments were performed. One group was stored at 4 °C. Other groups were placed in a thermal cycler, heated at 94°C for 3 min and cooled at 4°C for 5min (number of cycles 1, 2, 5, 10 and 100). The supernatant was collected from each condition. Protein concentration was measured by Micro BCA Protein Assay kit according to the manufacturer’s protocol (23235, Thermo Fisher). Solubilized cellular proteins from non-treated cells in LDS buffer were used as 100% in calculating eluted cellular proteins from the polyacrylamide beads.

#### Agarose Gel Electrophoresis

Agarose gel electrophoresis was performed through 2% agarose gel, E-Gel (G521802, Thermo Fisher Scientific) and DNA products were visualized by the imager LAS 4000 (FujiFilm).

#### Acquisition of Locational Information of Individual Antibodies on the Genome

Cell membranes and nuclear membranes were permeabilized using the following protocol. Single cells were treated with 1 x TBS containing 1% Triton X-100, 1 mM EDTA and 10% glycerol (TBS-Triton) for 15 min. The cells were then serially treated with 25%, 50%, 75% and 100% methanol for 5 min each, then treated with 75%, 50% and 25% methanol. The single cell was washed with the TBS-Triton.

In the “Annealing” step (**Fig. 1B**), the single cell was washed with “ThermoPol Mg (-) buffer” (20 mM Tri-HCl, pH8.8, 10 mM (NH4)2SO4, 10 mM KCl and 0.1% Triton X-100) and the supernatant was removed. ThermoPol Mg (-) buffer (15 μl/tube) containing 17.5 μM of the 1^st^ Random Primer (5’-/5Phos/CGACGCTNNNNNNNN-3’, Integrated DNA Technologies) was added to the single cell and incubated for 1 hour to deliver the random primer to the nucleus. The tube was incubated at 94 °C for 3 min and incubated on ice for 2 min or longer.

In the “Extension” step (**Fig. 1B**), a solution containing MgSO4 (final concentration 2 mM), NaCl (final concentration 300 mM) and dNTPs (final concentration 1.47 mM) was added followed by addition of the DNA polymerase DeepVent Exo(-) (1 unit, M0259L, New England BioLabs) and incubation for 4 hours on ice, 2 hours at 4°C, 2 hours at 10 °C, 2 hours at 20°C and 4 hours at 25°C.

In the “Antibody binding” step (**Fig. 1B**), the cell was washed and incubated with the ThermoPol Mg (-) buffer containing 300 mM NaCl, 10 mM EDTA and 1% BSA and 10% glycerol for 1 hour. Antibodies conjugated with the DNA probe were added (0.5 μg IgG/ml each) to the cell and incubated overnight on ice. The cell was washed with the ThermoPol Mg (-) buffer containing 300 mM NaCl.

In the “Proximity joining” and “Proximity ligation” steps (**Fig. 1B**), the cell was incubated with ThermoPol Mg (-) buffer containing 300 mM NaCl and 0.5 μM Ligation Adapter Probe (5’-AGCGTCGTGTAGGGAA-3’, Integrated DNA Technologies) for 1 hour at 25°C. The cell was washed once with 1x Quick Ligation Reaction buffer (M0202L, New England BioLabs). Quick Ligase (1 μl) and 1 x Quick Ligation Reaction buffer (19 μl) were added and incubated for 4 hours at 15 °C and 30 min at 25°C.

The ligated product of Antibody-DNA probe + 1^st^ Random primer was fully extended using the polymerase Bst 3.0 (M0374L, New England BioLabs). The cell was washed with ThermoPol Mg (-) buffer containing 300 mM NaCl. The polymerase solution was then added (320 unit/ml Bst 3.0 polymerase, 1.4 mM dNTPs and 2.5 mM MgSO4 in ThermoPol buffer). The cell was incubated for 4 hours at 4°C, 1 hour at 10°C, 1 hour at 20°C, 1 hour at 30°C, 1 hour at 40°C, 1 hour at 50°C and 1 hour at 65°C.

The extended products were amplified by a modified Multiple Annealing and Looping Based Amplification Cycles (MALBAC) (Lu et al. 2012; Zong et al. 2012). Two μM of the 2^nd^ Random primer (**Table S1**) was added and incubated for 2 hours. The cell and the extended products were heated at 94°C for 3 min and incubated on ice for 2 min. Bst 3.0 polymerase (1 l/tube) was added and incubated for 4 hours at 4°C, 30 min at 10°C, 30 min at 20°C, 30 min at 30°C, 30 min at 40°C, 30 min at 50°C, 60 min at 65°C and 3 min at 94°C. The supernatant was collected, and 20 l of buffer containing 10 mM Tris-HCl, pH 7.4, 0.1 mM EDTA and 0.05% Tween 20 was added. After overnight incubation at 4°C, the supernatant was collected; supernatant collection was repeated for a total of 3 times. The collected supernatant was combined into a 0.2 ml PCR tube. Four l of 10 x ThermoPol buffer (New England BioLabs) and dNTPs (total 1.4 mM) were added, and then the combined solution was heated at 94°C for 3 min and placed on ice for 2 min. Bst large fragment (2 μl/tube, M0275L, New England BioLabs) was added and incubated at 10°C for 45secs, 20°C for 45secs, 30°C for 45secs, 40°C for 45secs, 50°C for 45secs, 65°C for 2mins and 94°C for 20s. The tubes were then quickly quenched on ice. After quenching on ice, Bst large fragment (2 ul/tube) was added and then incubated at 10°C for 45secs, 20°C for 45secs, 30°C for 45secs, 40°C for 45secs, 50°C for 45secs, 65°C for 2mins and 94°C for 20s. The tubes were then quickly quenched on ice. The above cycles were repeated for a total of 8 times.

The products were further amplified by PCR using a primer (5’-ATCCATAGTGTCAGCAGGCT-3’) with DeepVent exo(-) polymerase (step1: 95°C-5 min, step2: 95°C-30 sec, step3: 60°C-30 sec, step4: 72°C-30 sec, step5: repeat step2-4 20 times, step6: 72°C-5 min and step 7: 4°C-forever). DNA was purified using UltraPure Phenol:Chloroform:Isoamyl Alcohol according to the manufacturer’s protocol (15593031, Thermo Fisher Scientific). Extra primers were removed by size-selective precipitation (Paithankar and Prasad 1991) using polyethylene glycol; 200 μg/tube linear acrylamide (AM9520, Thermo Fisher Scientific) and 2 M MgCl2 (final concentration 20 mM) were added, and then 50%(w/v) PEG8000 was added (final concentration 14%). After 20 min incubation, the tube was centrifuged at 1,3400 x g for 10 min. The pellet was washed with 80% ethanol 3 times and dried. After dissolving the pellet with 0.1x TE buffer, the amount of double-stranded DNA (dsDNA) was measured by Quant-iT PicoGreen dsDNA Assay kit (P11496, Thermo Fisher Scientific) and concentration of dsDNA was adjusted for *in vitro* transcription. The “reusable” single cell was stored at −20°C in TBS buffer containing 50% glycerol, 0.1% Triton X-100, 0.5% BSA and 1 mM EDTA) until the next round of experiments.

MALBAC, developed to amplify genomic DNA from single cells(Lu et al. 2012; Zong et al. 2012), can amplify all types of DNA. In our system, MALBAC could amplify desired products (antibody-DNA probe + 1^st^ random primer + genome sequence + 2^nd^ random primer) and genome-derived byproducts. Genome-derived byproducts were removed by the following steps. The antibody-DNA probe contains a T7 promoter sequence (**Table 1** and **Supplemental Fig. S2**). We converted the ligated products into RNA by *in vitro* transcription using HiScribe T7 High Yield RNA Synthesis Kit according to the manufacturer’s protocol (E2040S, New England BioLabs). DNA was digested using RNase-free DNase I according to the manufacturer’s protocol (EN0521, Thermo Fisher Scientific). RNA was purified using TRIzol LS Reagent according to the manufacturer’s protocol (10296028, Thermo Fisher Scientific). The purified RNA was converted into DNA using SuperScript IV Reverse Transcriptase according to the manufacturer’s protocol (18090010, Thermo Fisher Scientific) with a primer (5’-ATCCATAGTGTCAGCAGGCT-3’, RNase-free HPLC purified oligo, Integrated DNA Technologies) specific for the sequence in the 2^nd^ random primer (see “Reverse transcription” part in **Supplemental Fig. S2D**). Second strand synthesis was performed using the DNA polymerase, DeepVent Exo (-) (1 unit/50 μl) with a 3.68 M primer (5’-TAGCTAAGGTATCCTCCAGG-3’), 200 μM dNTPs and 1 x ThermoPol Reaction buffer. Extra sequences in the DNA products were removed by restriction-enzyme digestion with BciVI according to the manufacturer’s protocol (R0596L, New England BioLabs, see “Digestion with a restriction enzyme” in **Supplemental Fig. S2F**). Fragments larger than 49 bp were selected using E-gel 2% agarose (Thermo Fisher Scientific) and were extracted using Freeze ‘N Squeeze DNA Gel Extraction Spin Columns (Bio-Rad). The extracted DNA was purified using UltraPure Phenol:Chloroform:Isoamy Alcohol according to the manufacturer’s protocol (Thermo Fisher Scientific). The purified DNA was used for library construction using the Illumina TruSeq PCR free kit (Illumina) with TruSeq DNA index kit plate (96 samples, Illumina). Individual samples were labeled with unique Illumina indexes. The constructed libraries were sequenced by MiSeq, NextSeq 550 or NovaSeq 6000.

### Preparation of Reads before Mapping to the Genome

Illumina adapter sequences were trimmed using the trimming software TrimGalore (version 0.4.5, http://www.bioinformatics.babraham.ac.uk/projects/trim_galore/) with options [--paired --illumina - e 0.2 --length]. Trimmed reads were demultiplexed using antibody barcodes shown in Supplementary Table 2 by software FlexBar (version 3.0.3) (Dodt et al. 2012). Reads containing a reverse complement sequence of an antibody barcode were inverted using FastX tool kit (version 0.0.14, http://hannonlab.cshl.edu/fastx_toolkit/index.html). Left side sequence of reads from 5’ end to an antibody barcode-ligated sequence was trimmed by FlexBar. Right side sequence of reads from 3’ end to a cell barcode was also trimmed by FlexBar. After the trimming, an 8 nucleotide sequence of the 1^st^ random primer was removed and added to read ID as a unique molecular identifier by software UMI tools (version 1.0.1)(Smith et al. 2017). All codes used here are available at https://github.com/HidetakaOhnuki-NCI/REpi-seq/tree/master/001_Data_Prep. The outline of the Shell script is shown as a drawing on page 2 of the PDF file available at https://github.com/HidetakaOhnuki-NCI/REpi-seq/blob/master/000_Workflow_page1-11.pdf.

### Mapping to the Genome and Removing Amplification Duplicates

The trimmed reads are mapped to the human genome, GRCh38 using software Bowtie2 (version 2.3.5.1) (Langmead and Salzberg 2012). Amplification duplicates were removed by UMI tools based on the unique molecular identifier derived from 8 nt random sequences and mapped position on the genome. The unique mapped reads were used for the subsequent data analysis. All codes used here are available at https://github.com/HidetakaOhnuki-NCI/REpi-seq/tree/master/001_Data_Prep. Outline of the Shell script is shown as drawing on page 2 of the PDF file available at https://github.com/HidetakaOhnuki-NCI/REpi-seq/blob/master/000_Workflow_page1-11.pdf.

### Analysis of Locational Preservation of H3K27ac and H3K27me3 Marks over Repeated Experiments

The unique mapped reads of 8 single cells were combined. The combined datasets of the first experiment were converted into a BED file and used as a peak file in the next step. Signal distribution in genomic regions where signal was detected in the first experiment was analyzed by Homer (version 4.10.4) (Heinz et al. 2010). The signal distribution pattern was calculated by the Homer in the first, second and third repeated experiments. The output data were visualized as a line graph by Microsoft EXCEL. All codes used here are available at https://github.com/HidetakaOhnuki-NCI/REpi-seq/tree/master/002_Figure_2a-2b. Outline of the Shell script is shown as a drawing on page 3 of the PDF file available at https://github.com/HidetakaOhnuki-NCI/REpi-seq/blob/master/000_Workflow_page1-11.pdf

### Identification of Putative Signals without Data Aggregation from Other Single Cells

Datasets of the first, second and third repeated experiments were separately combined as reads derived from antibody or control IgG. Datasets of individual single cells remained as separate files. Randomized controls were generated from the combined antibody reads or control IgG using software BED tools (version 2.29.0) (Quinlan and Hall 2010). The randomized controls from each single cell were used to estimate stochastic background levels used in the following bootstrap statistical test. Genomic regions were subdivided into the 500 bp bin and the bin was shifted every 250 bp. Signals in the 500 bp bins were counted by BED tools with the option “intersect”. To estimate signal enrichment over control IgG, counts of control-IgG reads were subtracted from counts of antibody reads in each bin (*AbΔIgG*). To estimate accidental enrichment, counts of randomized control-IgG reads were subtracted from counts of randomized antibody reads (*Random AbΔRandom IgG*). The upper endpoint of the 99% confidence interval in the *Random AbΔRandom IgG* was calculated by bootstrap statistical test using R package “boot” (version 1.3-23). Regions with values of *AbΔIgG* greater than 99% confidence interval of *Random AbΔRandom IgG* were considered as signal-enriched regions compared to the background. The numbers of signal-enriched regions were counted and shown in **Figures 2C** and **2D**. Signals within the signal-enriched regions were counted (shown in **Fig.s 2D** and **2F**). These signals were considered as “putative” signals for further evaluation in the subsequent analyses. All codes used here are available at https://github.com/HidetakaOhnuki-NCI/REpi-seq/tree/master/003_Fiugre2c-2f. The outline of the Shell script, in combination with the R script, shown on page 4 of the PDF file available at https://github.com/HidetakaOhnuki-NCI/REpi-seq/blob/master/000_Workflow_page1-11.pdf.

### Identification of Putative Signals with Data Aggregation of Other Single Cells

For Med1 and 5hmC, the conventional approach was used to identify putative signals based on the commonality of signals among cells, using data aggregation of other single cells. The conventional approach is useful to increase the number of epigenetic marks, without repeating the experiments with single cells. The unique reads from 8 single cells were combined into one BAM file using SAM tools (version 1.9) (Li et al. 2009). Each randomized control was generated from reads of antibody or control IgG by BED tools. The number of reads in each bin was counted by BED tools with the option “intersect”. To estimate signal enrichment over control IgG, counts of control-IgG reads were subtracted from counts of antibody reads in each bin (*AbΔIgG*). To estimate accidental enrichment, counts of randomized control-IgG reads were subtracted from counts of randomized antibody reads (*Random AbΔRandom IgG*). The upper endpoint of 99% confidence interval for the *Random AbΔRandom IgG* was calculated by bootstrap statistical test using the R package “boot”. Regions with greater values in *AbΔIgG* than 99% confidence interval of *Random AbΔRandom IgG* were considered as signal-enriched regions compared to the background. The numbers of signal-enriched regions were counted and are shown in **Figures 2C** and **2D**. The numbers of signals in the significantly signal-enriched regions were counted and are shown in **Figures 2D** and **2F**. These signals were considered as “putative” signals for further evaluation in the subsequent analyses. All codes used here are available at https://github.com/HidetakaOhnuki-NCI/REpi-seq/tree/master/004_Figure_S4. Outline of the Shell script is shown on page 5 of the PDF file available at https://github.com/HidetakaOhnuki-NCI/REpi-seq/blob/master/000_Workflow_page1-11.pdf.

### Signal-Enriched Promoters and Enhancers

“Putative” H3K27ac signals identified by the bootstrap test were used to determine signal-enriched promoters and enhancers. A list of human promoters was downloaded from Eukaryotic Promoter Database (Dreos et al. 2017) of the Swiss Institute of Bioinformatics as a <u>BED file</u> (version 006, assembly: hg38). A list of human enhancers was downloaded from the Human ACtive Enhancers to interpret Regulatory variants (HACER) atlas (Wang et al. 2019). To compare the results of REpi-seq with data from bulk cell analysis, bulk ChIP-seq dataset of K562 cells was used (from ENCODE, Data set ID: ENCFF301TVL). To estimate the frequency of accidental detection, the “putative” H3K27ac signals in REpi-seq and bulk ChIP-seq were randomized using BED tools, and the generated data were used as controls. Numbers of putative H3K27ac signals and random control in promoters and enhancers ware counted by BED tools with the option “intersect”. Signal counts were normalized based on the length of promoters or enhancers (per kilobase) and the number of input signals (per million input). Bootstrap statistical test was performed for the random controls to determine the upper endpoint of the 95% confidence interval. The value of the 95% confidence interval was used to select significantly signal-enriched promoters and enhancers. The number of significantly signal-enriched enhancers and promoters was counted for bulk ChIP-seq and REpi-seq data. All codes used here are available at https://github.com/HidetakaOhnuki-NCI/REpi-seq/tree/master/005_Figure3a-3d. The workflow and data flow of the Shell script is shown on page 6 of the PDF file available at https://github.com/HidetakaOhnuki-NCI/REpi-seq/blob/master/000_Workflow_page1-11.pdf.

### Enhancer Classification

Enhancers in the HACER datasets were classified based on relative ratios Log2(H3K27ac/H3K27me3). “Putative” signals of H3K27ac, H3K27me3, Med1 and 5hmC in enhancers were counted using BED tools with option “map”. Signal counts were normalized based on the length of enhancers (per kilobase) and based on the number of input signals (per million input). A BED file of enhancers was split based on Log2(H3K27ac/H3K27me3). Average signal counts were calculated in the classified enhancers. All codes used here are available at https://github.com/HidetakaOhnuki-NCI/REpi-seq/tree/master/006_Figure3e-3h. The workflow and data flow of the Shell script was shown on page 7 of the PDF file available at https://github.com/HidetakaOhnuki-NCI/REpi-seq/blob/master/000_Workflow_page1-11.pdf.

### Genes Interacting with Active Enhancers

Active enhancers were classified based on relative ratios Log2(H3K27ac/H3K27me3) calculated using “putative” signals with a p-value < 0.01. To compare the results of REpi-seq with results of bulk ChIP-seq, H3K27ac and H3K27me3 data sets of bulk ChIP-seq were used (ENCODE, dataset ID: ENCFF301TVL and ENCFF330YFF). The number of putative H3K27ac or H3K27me3 signals in enhancers was counted by BED tools. Signal counts were normalized based on the length of enhancers (per kilobase) and based on the number of input signals (per million input). Relative ratio Log2(H3K27ac/H3K27me3) was calculated from the normalized signal counts, and enhancers having a value greater than 0 were selected as putative active enhancers. The putative active enhancers were separated based on cell-type specificity defined by the Human ACtive Enhancers to interpret Regulatory variants (HACER) atlas (Wang et al. 2019). A list of genes, which interact with the identified putative active enhancers was generated based on experimentally validated information in the HACER. The number of genes was counted; overlapping genes between bulk ChIP-seq and REpi-seq are shown in **Figures 4A** and **4B**. All codes used are available at https://github.com/HidetakaOhnuki-NCI/REpi-seq/tree/master/007_Figure4a-4b. The workflow and data flow of the Shell script is shown on page 8 of the PDF file available at https://github.com/HidetakaOhnuki-NCI/REpi-seq/blob/master/000_Workflow_page1-11.pdf.

### Pathway Enrichment Analysis

Functions of genes proximal to the putative active enhancers were analyzed by Pathway Enrichment Analysis (Kramer et al. 2014). A BED file was generated by combining BED files of enhancers (HACER) and promoters (Eukaryotic Promoter Database of the Swiss Institute of Bioinformatics). H3K27ac and H3K27me3 signals from REpi-seq and bulk ChIP-seq (ENCFF301TVL and ENCFF330YFF) in enhancers and promoters were counted by BED tools. Signal counts ware normalized based on the length of enhancers (per kilobase) and the number of input signals (per million input). Relative ratio Log2(H3K27ac/H3K27me3) was calculated from the normalized signal counts. A list of genes was generated form HACER and the promoter datasets. The epigenetic score of a gene was calculated by the sum of epigenetic scores of proximal enhancers and promoters. The epigenetic score of genes was used as input data for Ingenuity Pathway Analysis (Qiagen). For RNA-seq data from bulk K562 and H1 cells, datasets were downloaded from the Cancer Cell Line Encyclopedia (Broad Institute) and ENCODE (dataset ID: ENCFF093NEQ). The number of reads per kilobase per million (RPKM) was converted to Log2, and the difference between K562 and H1 cells were calculated. The difference was converted into a ranking metric, and the data were converted into a text file. The text file was used as input data for Ingenuity Pathway Analysis. The top 5,000 transcripts from K562 and H1 cells were used in the Pathway Enrichment Analysis. All codes used are available at https://github.com/HidetakaOhnuki-NCI/REpi-seq/tree/master/008_Figure4d. The workflow and data flow of the Shell script is shown on page 9 of the PDF file available at https://github.com/HidetakaOhnuki-NCI/REpi-seq/blob/master/000_Workflow_page1-11.pdf.

### t-Stochastic Neighbor Embedding (t-SNE)

The t-SNE plot was generated from the epigenetic status of enhancers and promoters. Bulk ChIP-seq data of H3K27ac and H3K27me3 were downloaded from ENCODE (258 datasets, the list is shown in Table S4). Signals from the bulk ChIP-seq in enhancers (HACER) and promoters (Eukaryotic Promoter Database of the Swiss Institute of Bioinformatics) were counted by BED tools. Putative signals of each single cell were counted by BED tools. Counts were normalized based on the length of enhancers or promoters (per kilobase) and based on the input number (per million inputs). Relative ratios Log2(H3K27ac/H3K27me3) were calculated from the normalized signal counts. The Log2(H3K27ac/H3K27me3) values of enhancers and promoters were used as input data for software SeqGeq (FlowJo) and t-SNE plot was generated. All codes are available at https://github.com/HidetakaOhnuki-NCI/REpi-seq/tree/master/009_Figure5a. The workflow and data flow of the Shell script was shown on page 10 of the PDF file available at https://github.com/HidetakaOhnuki-NCI/REpi-seq/blob/master/000_Workflow_page1-11.pdf.

### Visualization of K562-Cell-Type-Specific and Non-Specific, Active Enhancers with Med1 and 5hmC Signals

Putative signals of Med1 and 5hmC in individual cells were extracted from unique reads using BED files generated in **Supplemental Figure S4** by BED tools. The extracted reads (BAM files) of individual cells were converted into BED files by BED tools with option “bamtobed”. Signal-enriched regions and signals of H3K27ac and H3K27me3 identified by bootstrap test (P < 0.01) were used to calculate relative ratios of Log2(H3K27ac/H3K27me3) in each single cell. Active enhancers (P < 0.05) compared to random controls were identified by a bootstrap statistical test in each single cell. Datasets of active enhancers were converted into TDF files by software IGVtools (version 2.7.2) (Robinson et al. 2011). The TDF files and BED files were visualized by Integrated Genome Viewer, and data are shown in **Figure 5B-5E** and **Supplemental Figure S5A-S5C**. All codes used here are available at https://github.com/HidetakaOhnuki-NCI/REpi-seq/tree/master/010_Figure5b-5e%2BFigureS5a-S5c. Workflow and data flow of the Shell script is shown on page 10 of the PDF file available at https://github.com/HidetakaOhnuki-NCI/REpi-seq/blob/master/000_Workflow_page1-11.pdf.

### General Reagents

In the experiments above, the following reagents were used; 10 x TBS, pH7.4 (351-086-101, Quality Biological), glycerol (15514-011, Thermo Fisher Scientific), 0.5 M EDTA, pH8.0 (15575-038, Thermo Fisher), BSA (B6917-100MG, Sigma-Aldrich), 10 x PBS, pH7.4 (70011-044, Thermo Fisher Scientific), Tween 20 (P1379-500ML, Sigma-Aldrich), 0.2 ml PCR tubes (A30588, Thermo Fisher Scientific), Tri-HCl, pH8.8 (T1588, Teknova), Ammonium sulfate (A4418-100G, Sigma-Aldrich), 2 M KCl (AM9640G, Thermo Fisher Scientific), Triton X-100 (T8787-50ML, Sigma-Aldrich), MgSO4(M3409-10×1ML, Sigma-Aldrich), 5 M NaCl (AM9760G, Thermo Fisher Scientific), 10 mM dNTP mix (R0192, Thermo Fisher Scientific), polyethylene glycol 8000 (P5413-500G, Sigma-Aldrich), 2 M MgCl2 (340-034-721, Quality Biological), TE buffer (351-010-131, Quality Biological) and UltraPure Water (10977-015, Thermo Fisher Scientific).

---

**Supplemental Figure S1.**
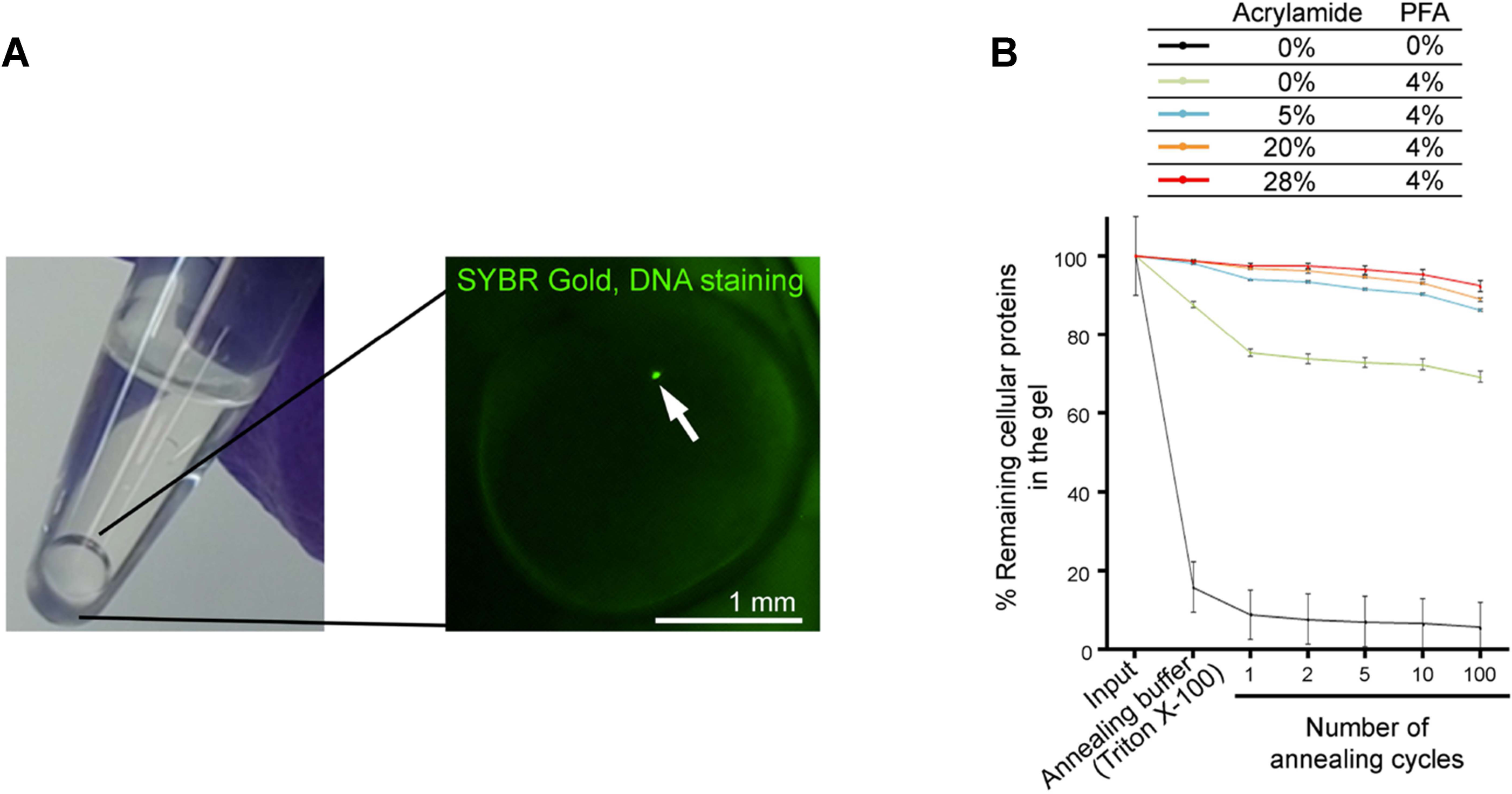
Generating “Reusable” Single Cells, Related to Figure 1 (A) A “reusable” single cell in a PCR tube. A single cell was embedded and anchored to a polyacrylamide (4%) scaffold. DNA in the cell nucleus was visualized by SYBR Gold. (B) Preservation of cellular proteins during heating and cooling cycles. Preservation of cellular proteins in non­ fixed (0% acrylamide and 0% PFA), PFA-only-fixed (0% acrylamide and 4% PFA) and reusable (5-28% acrylamide and 4% PFA) single cells after heating and cooling cycles. The cells (3.33 x 105 cells/tube) were mixed with PBS containing 3.88% acrylamide, 0.12% bis-acrylamide and 1% ammonium persulfate. The cells were embedded into polyacrylamide by adding mineral oil containing 0.2% N,N,N’,N’-tetramethylethane-1,2-diamine. Eluted proteins were measured by Micro BCA after the heating and cooling cycles (94 °C for 3 min and 4 °C for 10 min). Inputs are total proteins from 3.33 x 105 cells. Each data point represents the mean of triplicate experiments. Error bars reflect standard deviations.

**Supplemental Figure S2.**
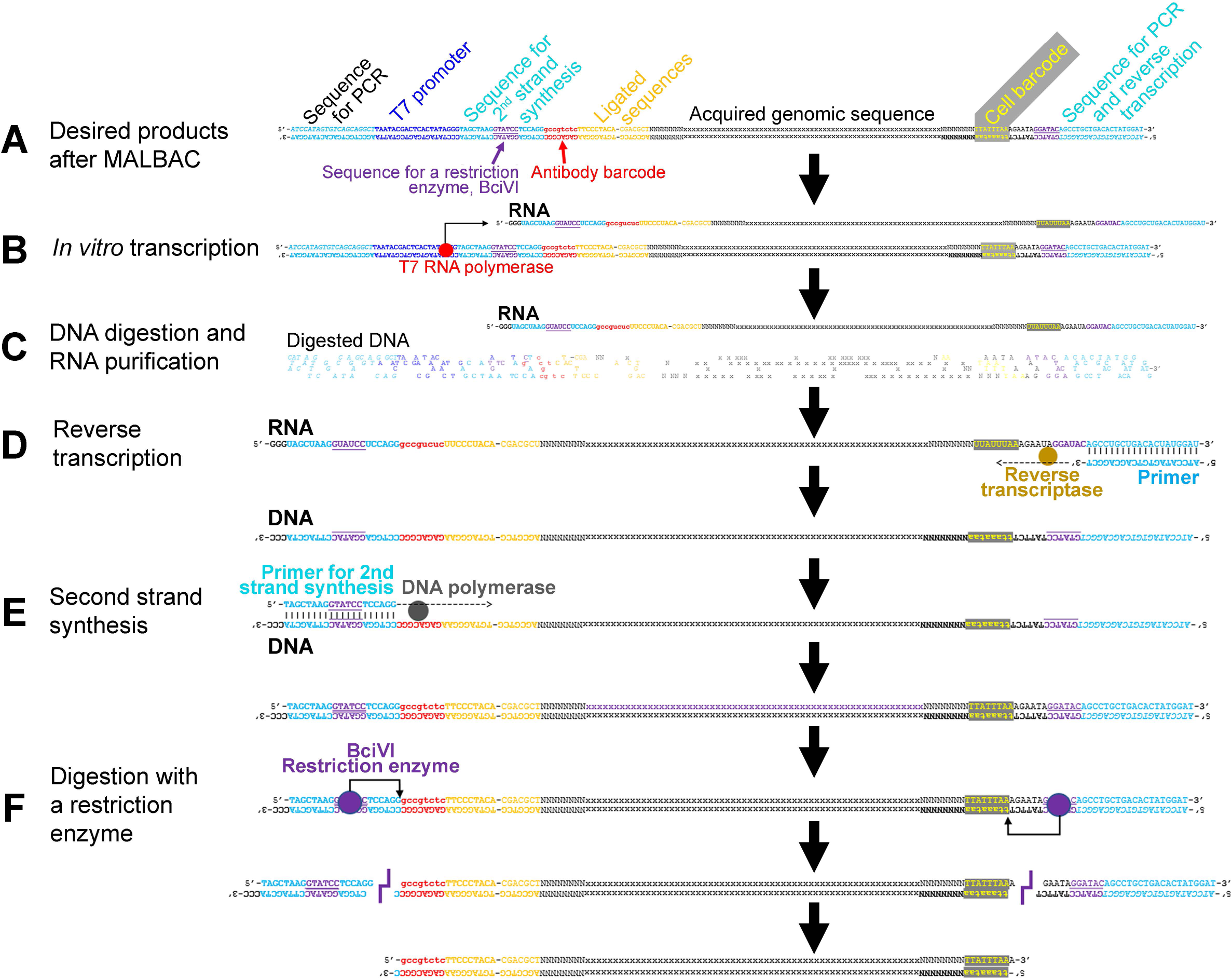
Representation of a Method to Reduce Genome-Derived Byproducts Generated by MALBAC in the Method, Related to Figure 1 (A) Desired products from MALBAC at the end of Figure 1b. (B) Conversion of the desired products into RNA by in vitro transcription using T7 promoter in the antibody probe. (C) Digestion of DNA including genome-derived byproducts of MALBAC, and RNA purification by phase separation. (D) Reverse transcription of the purified RNA (E) Second strand synthesis using a specific primer. (F) Removal of extra sequences by digestion with the restriction enzyme BciVL

**Supplemental Figure S3.**
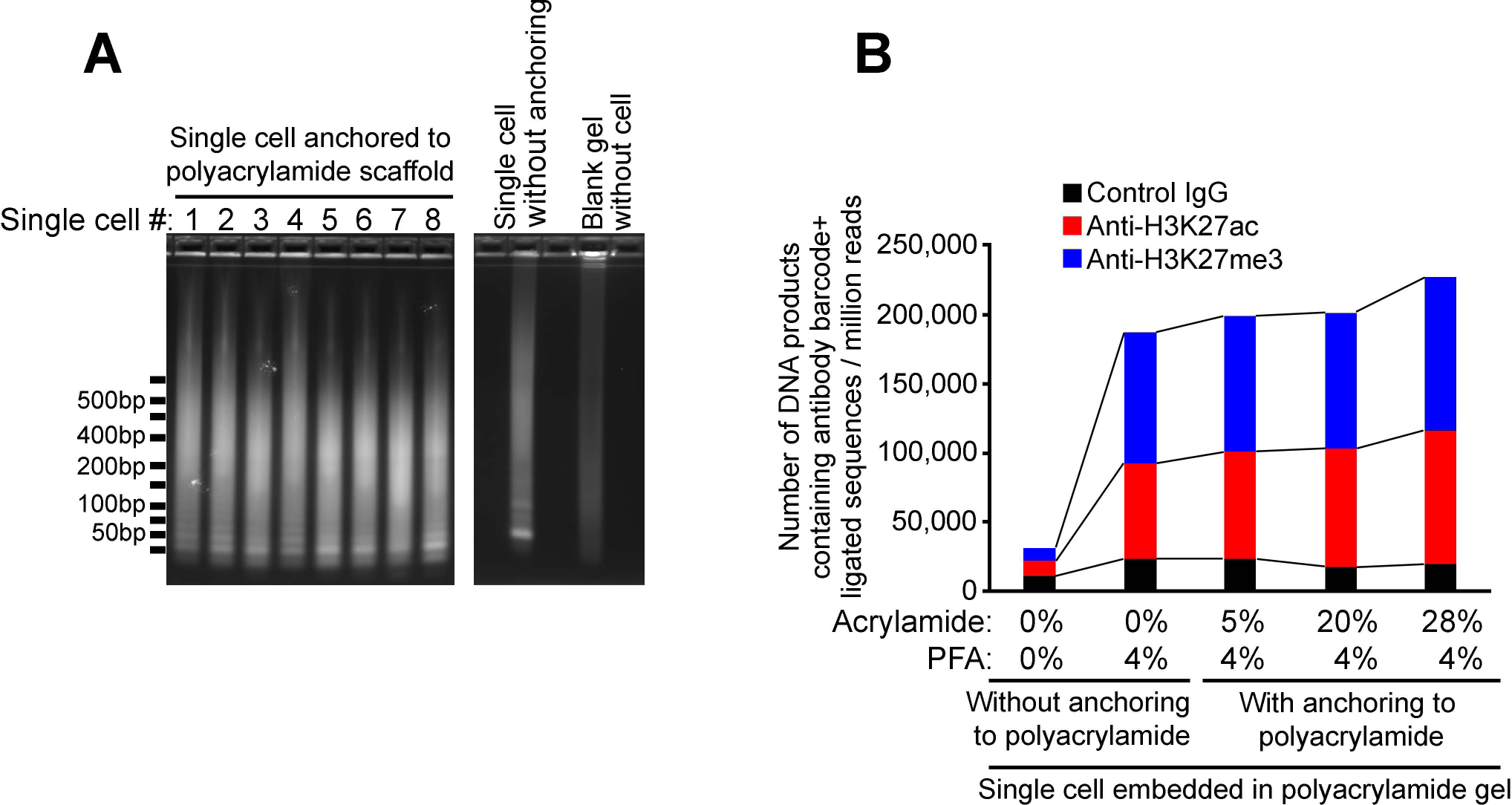
DNA Products from Single Cells Embedded in a Polyacrylamide Gel, Related to Figure 1 (A) Gel images of products at the end of the series of reactions. DNA was visualized with SYBR Gold in 2% agarose gels. (B) Numbers of DNA products containing sequences of the antibody probe and the first random primer. Single cells were embedded in a 4% polyacrylamide gel bead with or without anchoring the proteins to the polyacrylamide scaffold.

**Supplemental Figure S4.**
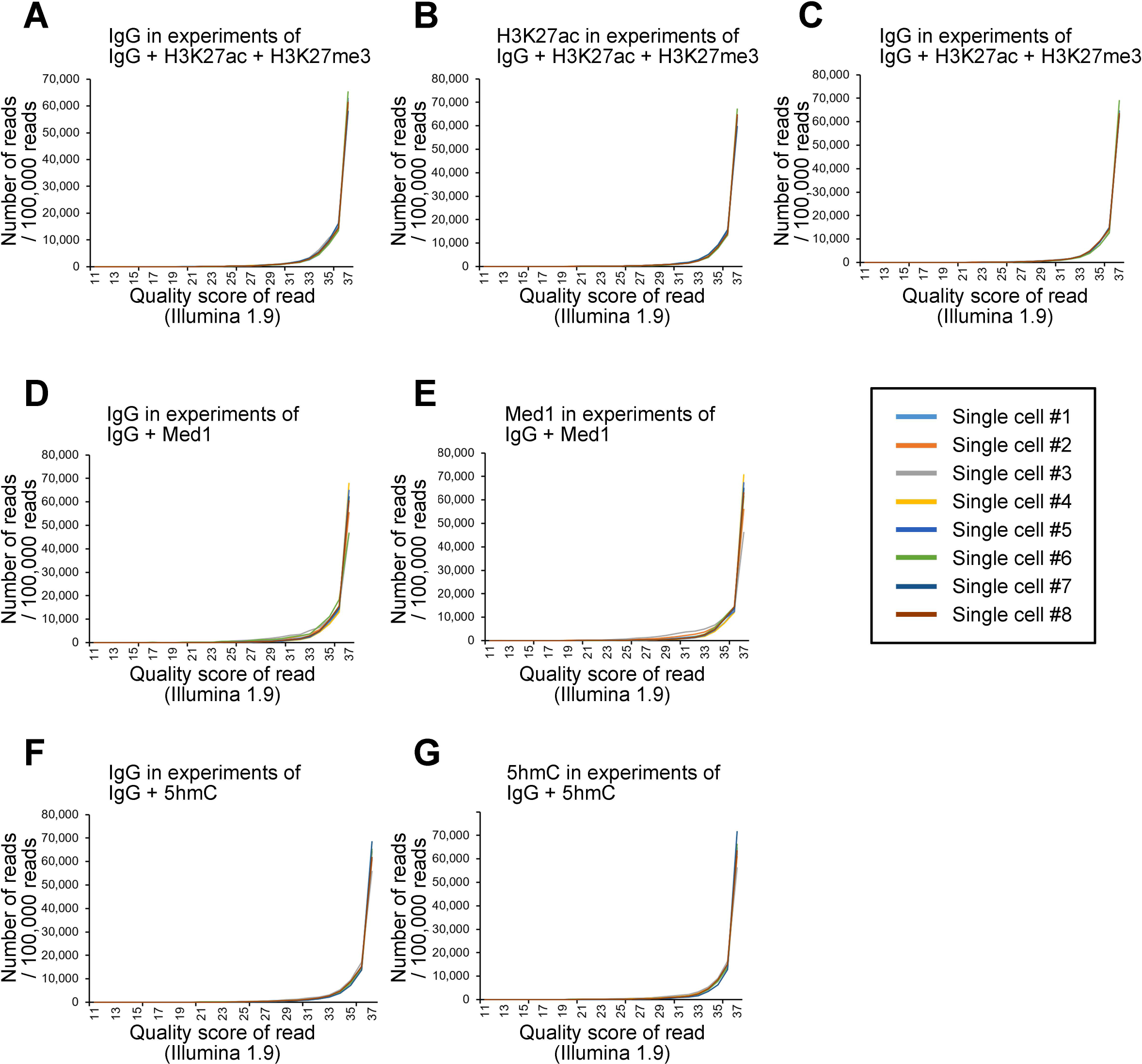
Frequency of sequencing quality score in mapped reads (A-C) Frequency of sequencing quality scores in mapped reads derived from control lgG (A), anti-H3K27ac (B) and anti-H3K27me3 (C). Control lgG, anti-H3K27ac and anti-H3K27me3 were mixed and reacted with a single cell. (D-E) Frequency of sequencing quality scores in mapped reads derived from control lgG (D) and anti-Med1 (E). Control lgG and anti-Med1 were mixed and reacted with a single cell. (F-G) Frequency of sequencing quality scores in mapped reads derived from control lgG (F) and anti-5hmC (G). Control lgG and anti-5hmC were mixed and reacted with a single cell. The generated products were sequenced and mapped to the genome. The probability of an sequence error in the mapped reads is shown as a quality score of a read (lllumina 1.9).

**Supplemental Figure S5.**
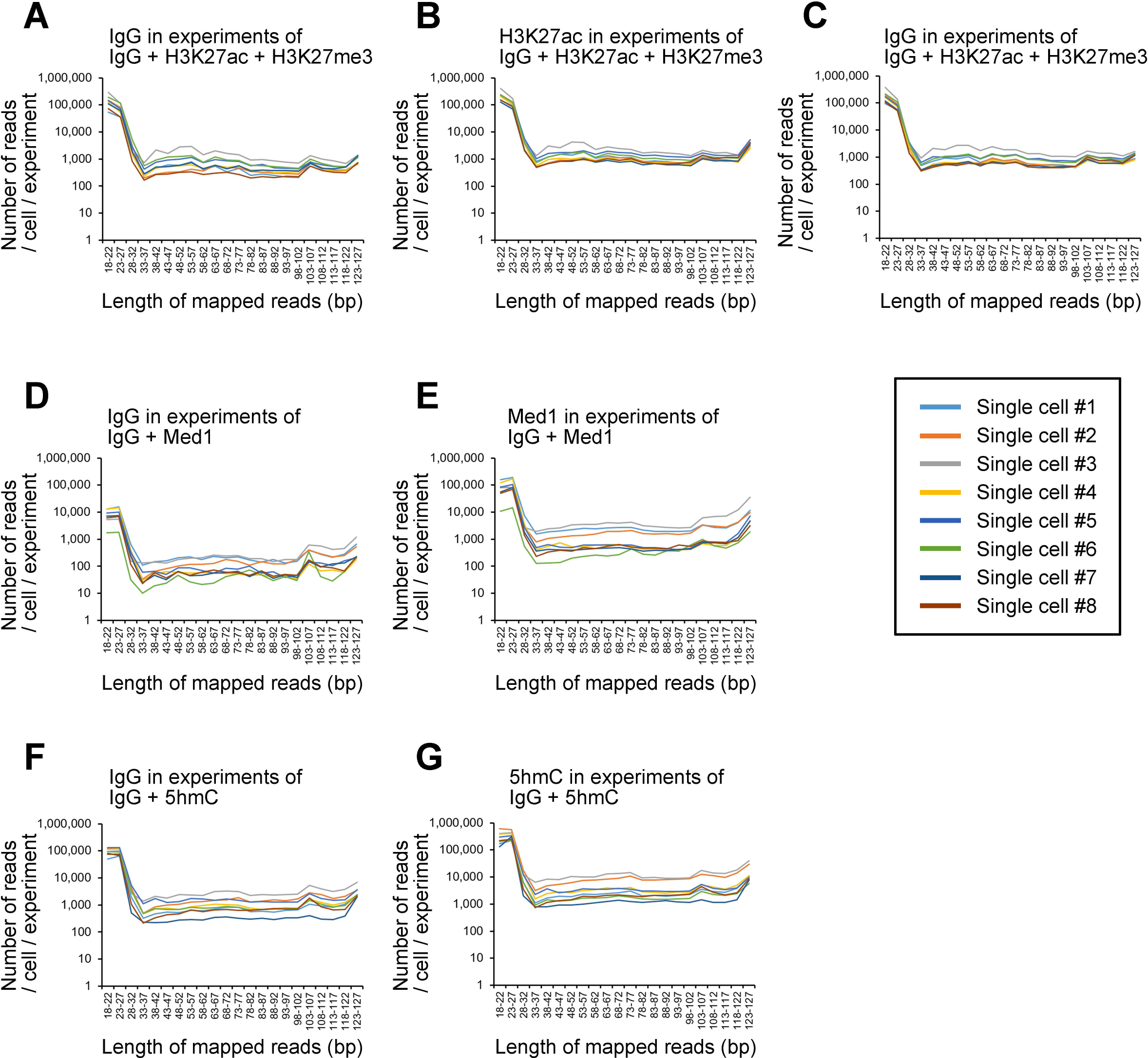
Length of added genomic sequences to the random primer of ligated products in mapped reads of a single cell. (A-C) Length of added genomic sequences to the random primer of ligated products in mapped reads derived from control lgG (A), anti-H3K27ac (B) and anti-H3K27me3 (C). Control lgG, anti-H3K27ac and anti-H3K27me3 were mixed and reacted with a single cell. (D-E) Length of added genomic sequences to the random primer of ligated products in mapped reads derived from control lgG (D) and anti­ Med1 (E).Control lgG and anti-Med1 were mixed and reacted with a single cell. (F-G) Length of added genomic sequences to the random primer of ligated products in mapped reads derived from control lgG (F) and anti-5hmC (G). Control lgG and anti-5hmC were mixed and reacted with a single cell. The generated products were sequenced and mapped to the genome. Each line indicates number of reads per cell in one experiment.

**Supplemental Figure S6.**
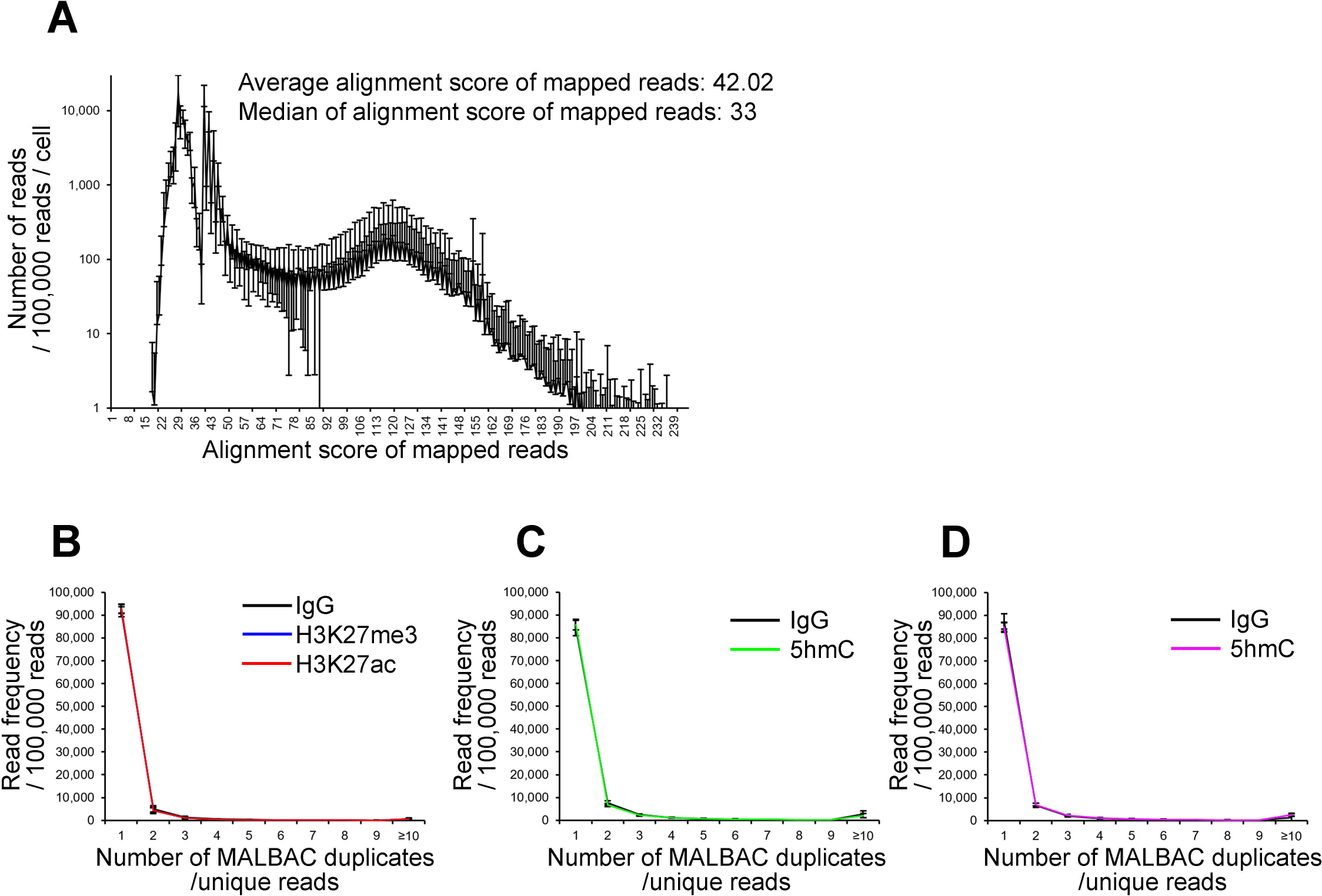
Mapping Quality Scores and Duplication Rates by MALBAC (A) Frequency of mapping quality score in mapped reads to the genome. (B-D) Frequency of duplication by MALBAC in lgG + H3K27me3 + H3K27ac (B), lgG + 5hmC (C) and lgG + HP1g (D). Error bar is standard diviation.

**Supplemental Figure 57.**
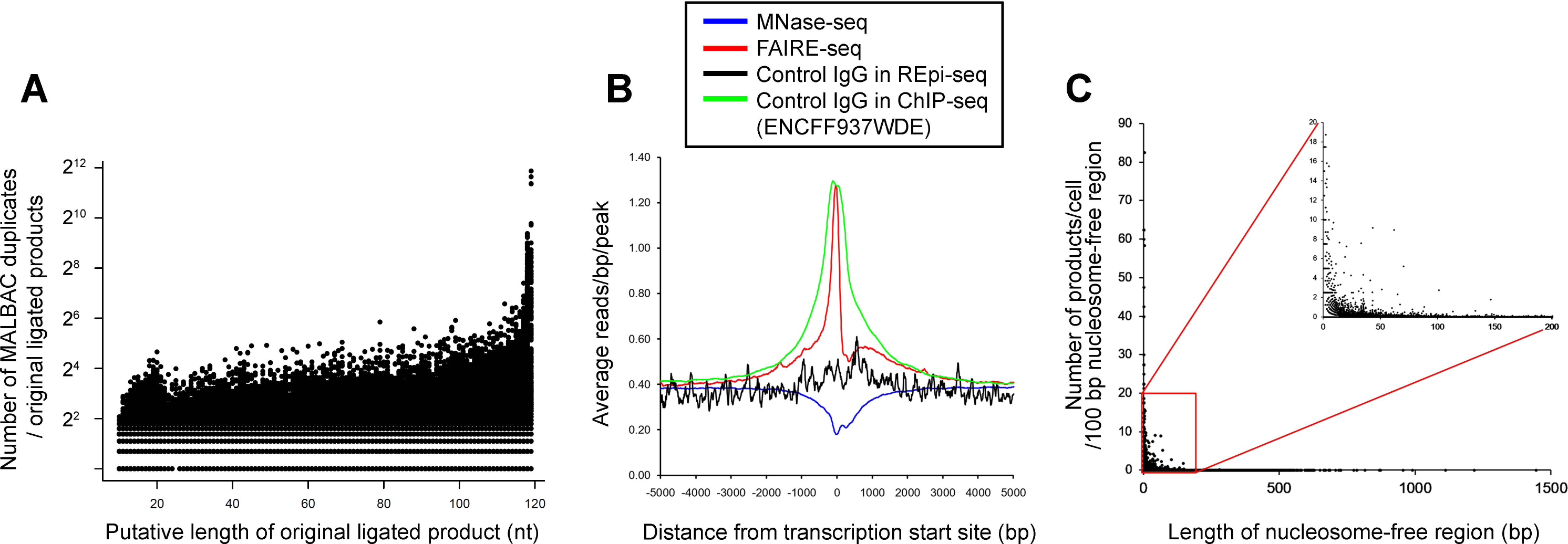
Evaluation of bias in nucleosome-free regions (A) Long ligated products generate higher numbers of MALBAC duplicates. MALBAC duplicates were identified and counted using combinations of antibody barcodes, 8 nucleotides random sequences in the first random primer and acquired genomic sequences. The longest of each ligated products, used as a putative original ligated product, served as a template in MALBAC (shown in X-axis). (B) Control lgG reads are less frequent in nucleosome-free regions at transcription start sites. (C) Inverse correlation between length of nucleosome-free regions and number of MALBAC duplicates.

**Supplemental Figure S8.**
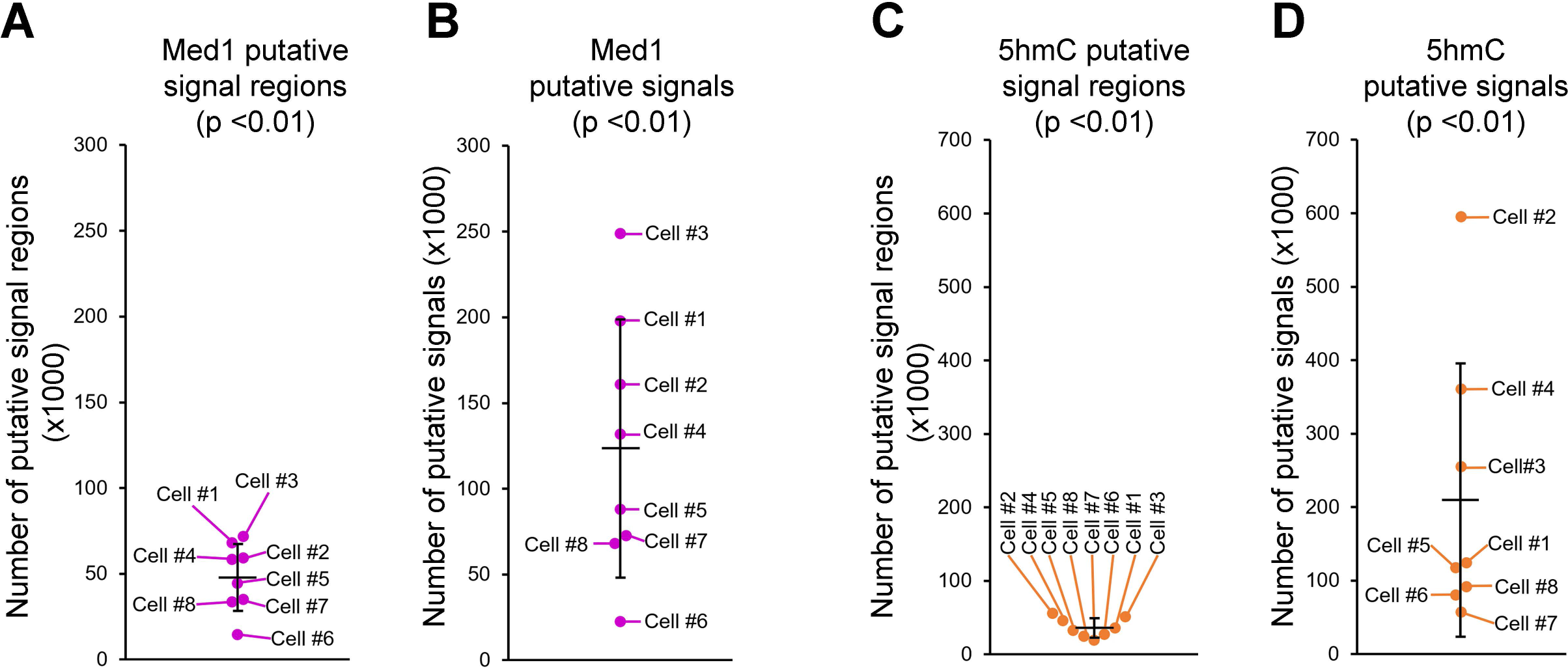
Reanalyzing the Same Single Cells with Different Antibodies, Related to Figure 3G and 3H (A and C) Signal-enriched genomic regions of mediator complex subunit 1 (Med1, A) and 5-hydroxymethylcytosine (5hmC, C)in a cell. Signal enriched regions were determined by 2 criteria. The first criterion is signal enrichment over control lgG in a genomic region (500 bp bin size). The second criterion is statistical significance compared to random controls of antibody and control lgG (see STAR Method for the detail). Genomic regions in the upper 1% of the distribution were considered as putative signal regions. (B and D) Numbers of putative signals per cell from Med1 (B) and 5hmC (D). Numbers of putative signals in the signal-enriched genomic regions were counted in every single cell.

**Supplemental Figure S9.**
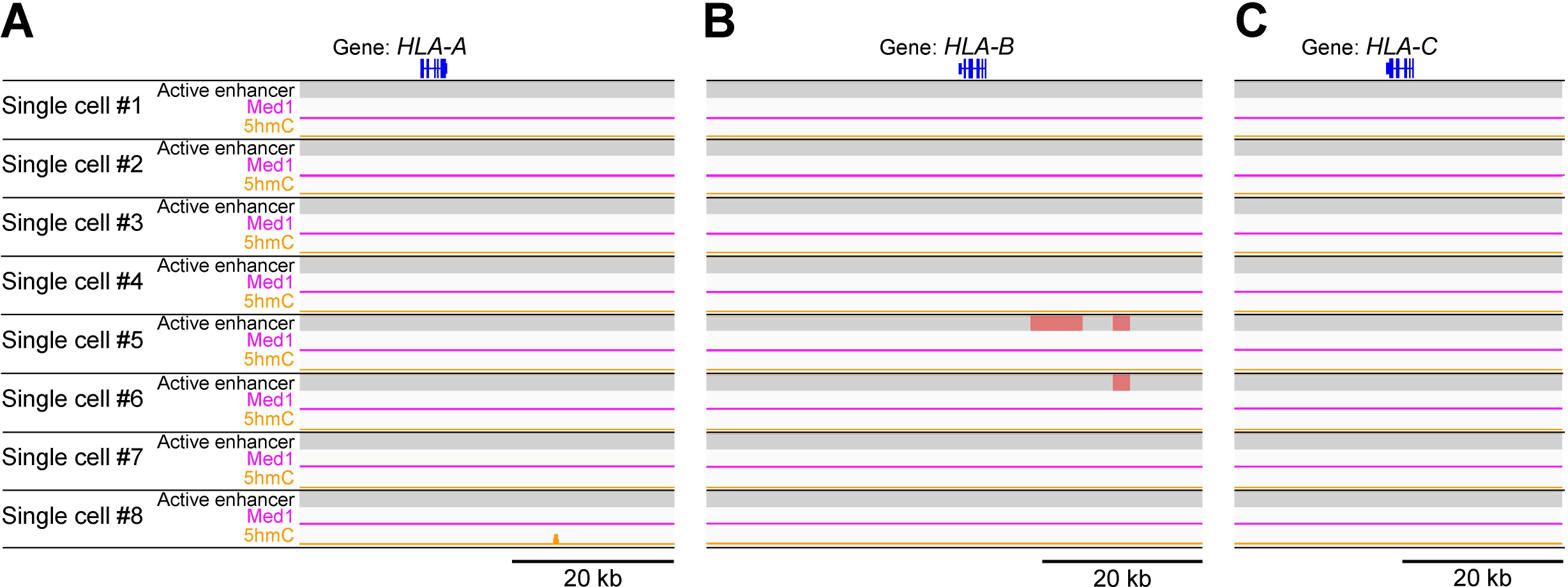
Epigenetic Status of Known Inactive Loci HLA-A, HLA-B and HLA-C in K562 Single Cells, Related to Figure 5 (A-C) Active enhancers [Log2(H3K27ac/H3K27me3),]Med1 and 5hmC in known inactive loci HLA-A, HLA-8 and HLA-C. Signals of H3K27ac, H3K27me3, Med1 and 5hmC (P < 0.01) were used as input data. Active enhancers were identified based on significant H3K27ac dominance in relative ratio Log2(H3K27ac/H3K27me3) compared to randomized controls of H3K27ac and H3K27me3 (P < 0.05, bootstrap test). Known K562-cell-type-specific, active enhancers of K562 cells were not detected. K562-cell-type non-specific, active enhancers (red) are active enhancers identified in other cells, but not in bulk K562 cells.

**Supplemental Table S1.**
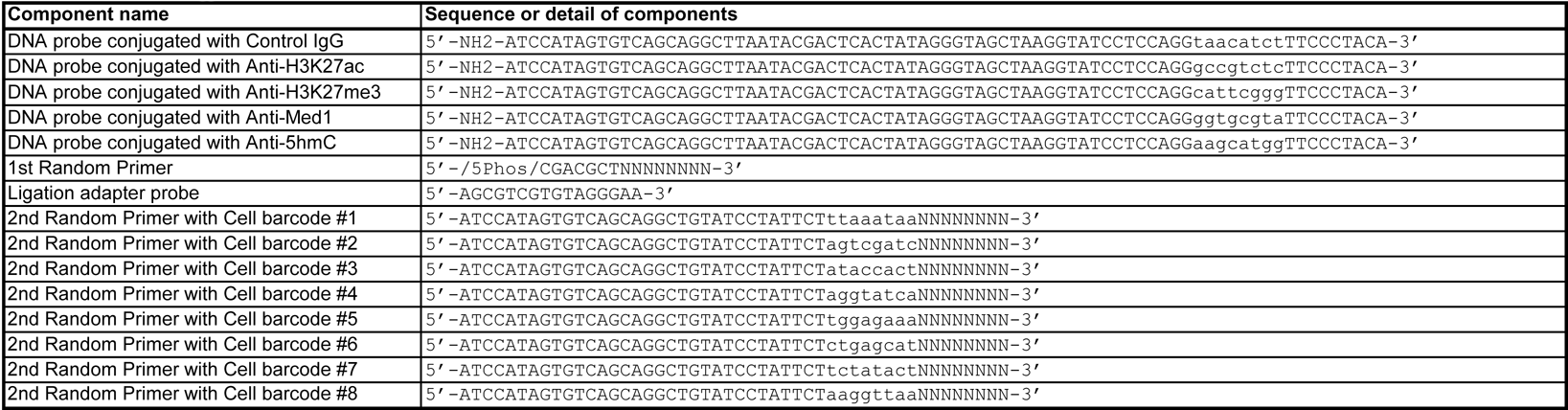
DNA components of the method

**Supplemental Table S2.** Sequences of antibody barcodes

| Antibody name | Forward sequence | Reverse complement sequence |
| --- | --- | --- |
| H3K27me3 | CATTCGGG | CCCGAATG |
| H3K27ac | GCCGTCTC | GAGACGGC |
| Med1 | GGTGCGTA | TACGCACC |
| 5hmC | AAGCATGG | CCATGCTT |
| Control IgG | TAACATCT | AGATGTTA |

**Supplemental Table S3.**
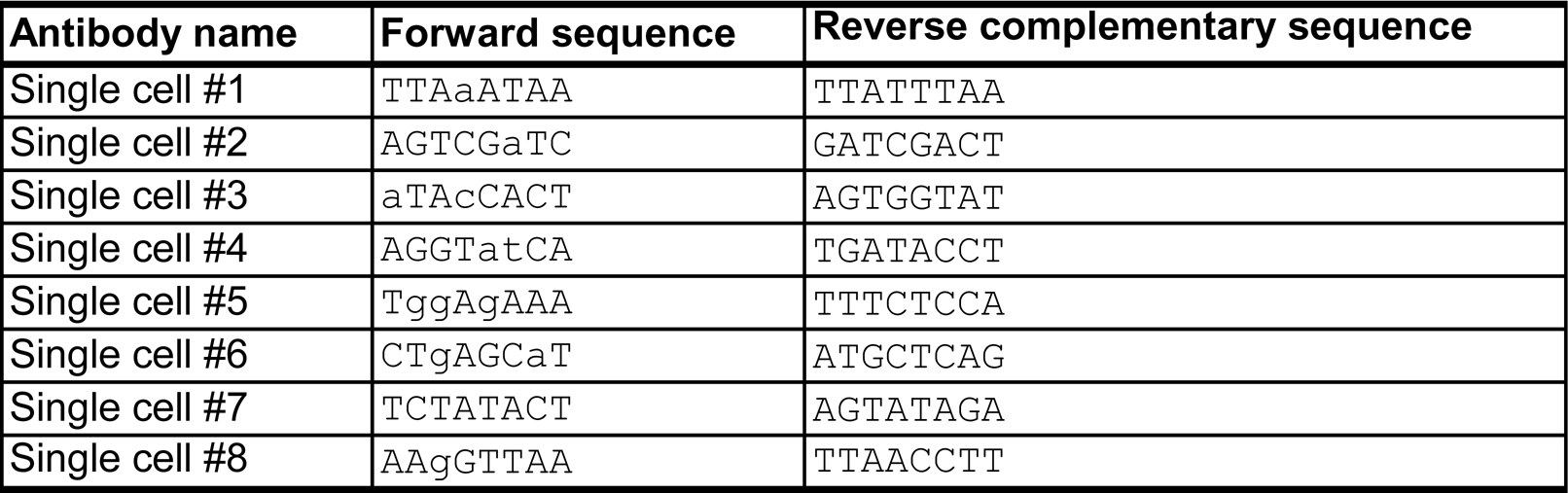
Sequences of cell barcodes

| Antibody name | Forward sequence | Reverse complementary sequence |
| --- | --- | --- |
| Single cell #1 | TTAaATAA | TTATTTAA |
| Single cell #2 | AGTCGaTC | GATCGACT |
| Single cell #3 | aTAcCACT | AGTGGTAT |
| Single cell #4 | AGGTatCA | TGATACCT |
| Single cell #5 | TggAgAAA | TTTCTCCA |
| Single cell #6 | CTgAGCaT | ATGCTCAG |
| Single cell #7 | TCTATACT | AGTATAGA |
| Single cell #8 | AAgGTTAA | TTAACCTT |

**Supplemental Table S4.**
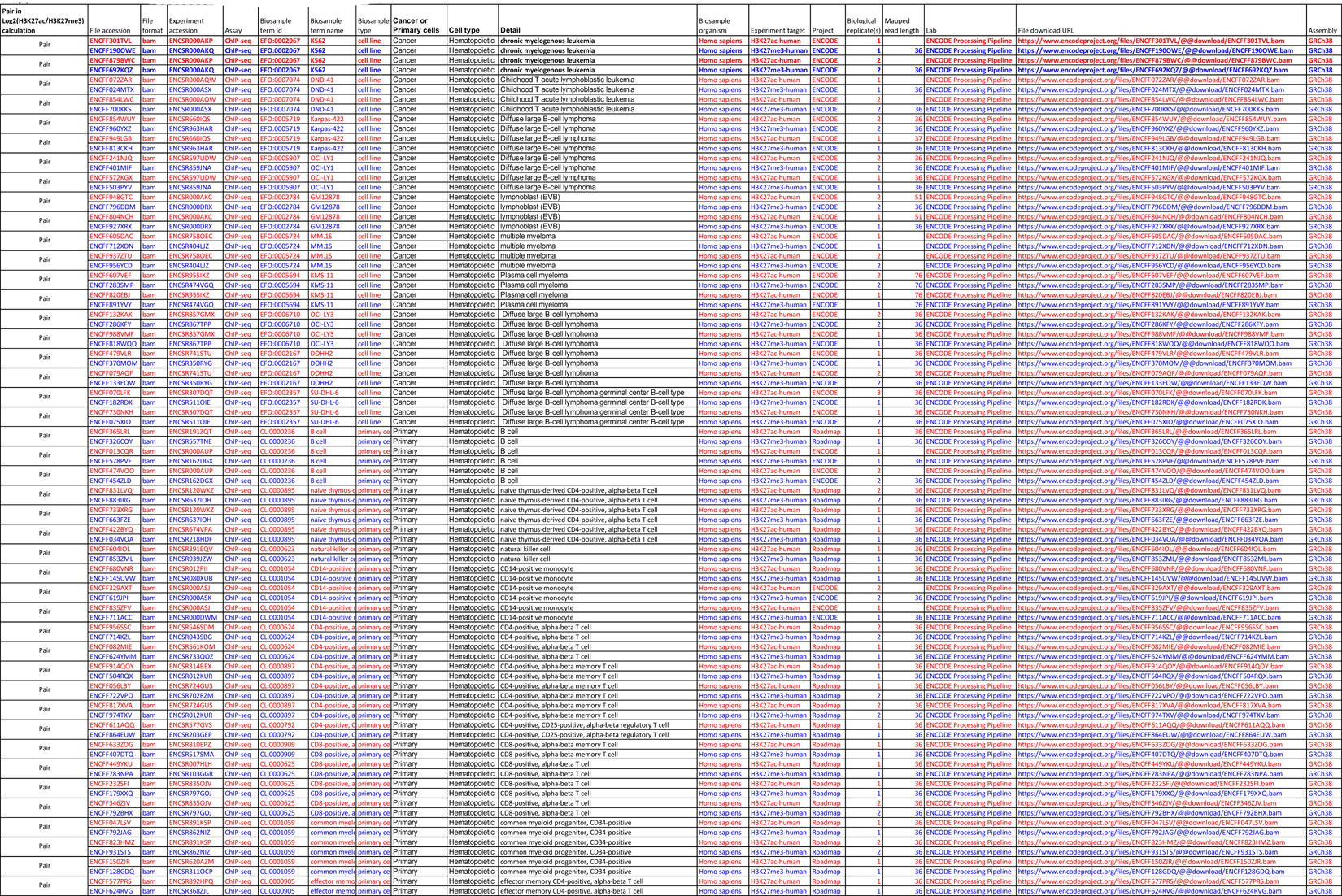
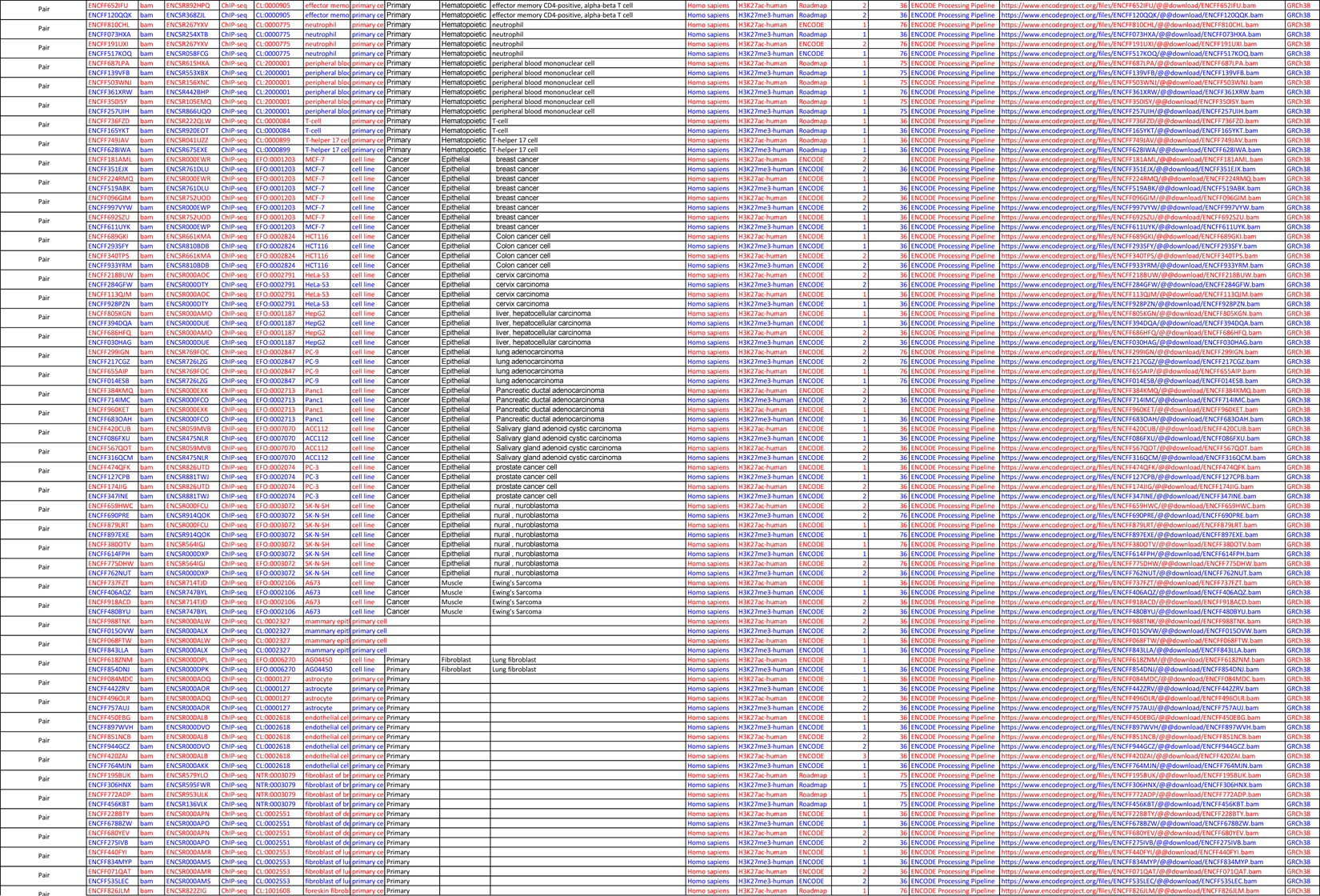
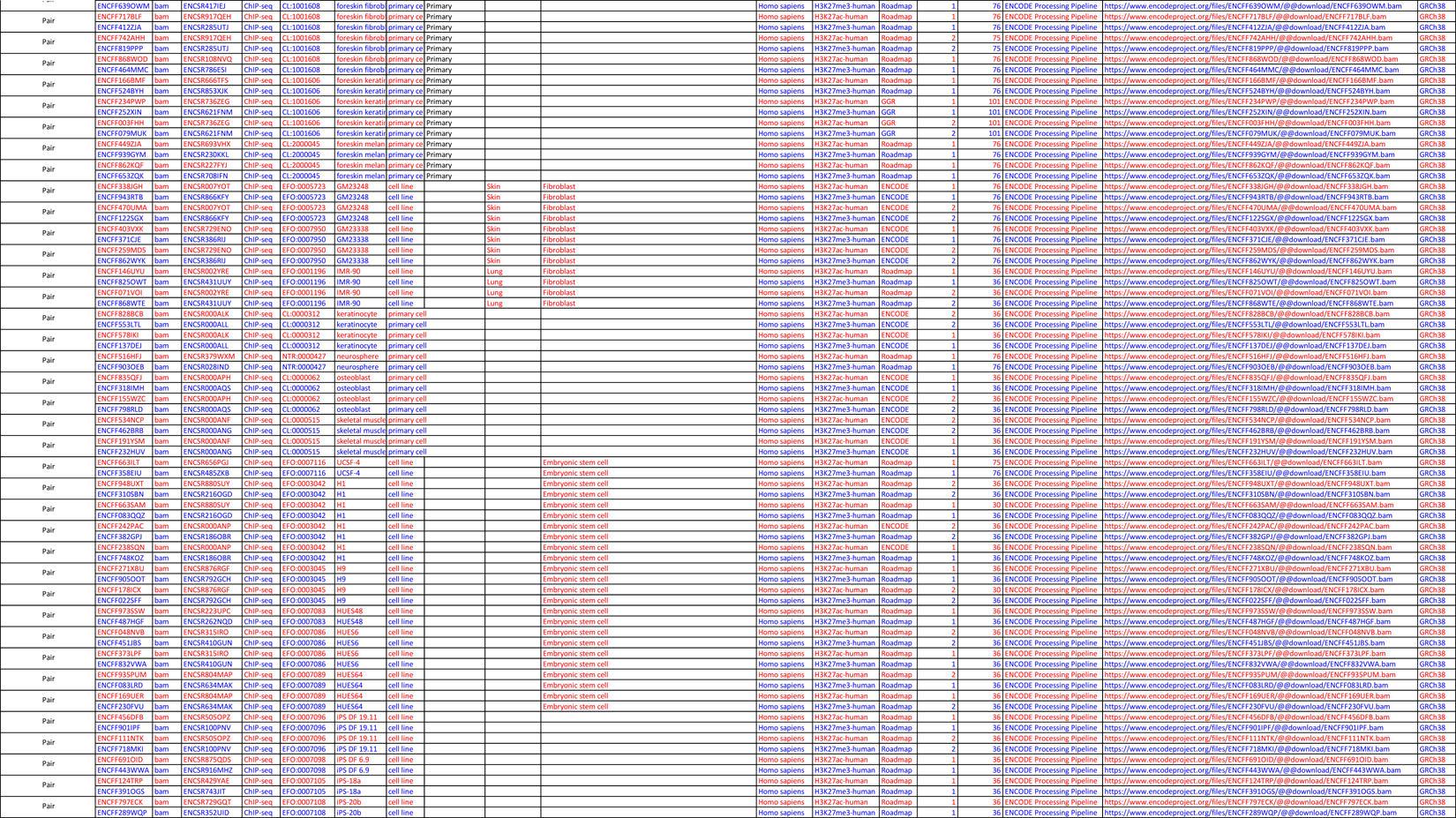
Datasets of bulk ChIP-seq used for t-SNE plots

## References

1. Aibar S, Gonzalez-Blas CB, Moerman T, Huynh-Thu VA, Imrichova H, Hulselmans G, Rambow F, Marine JC, Geurts P, Aerts J et al. 2017. SCENIC: single-cell regulatory network inference and clustering. Nat Methods 14: 1083–1086.

2. Andersson LC, Nilsson K, Gahmberg CG. 1979. K562--a human erythroleukemic cell line. Int J Cancer 23: 143–147.

3. Bock C, Kiskinis E, Verstappen G, Gu H, Boulting G, Smith ZD, Ziller M, Croft GF, Amoroso MW, Oakley DH et al. 2011. Reference Maps of human ES and iPS cell variation enable high-throughput characterization of pluripotent cell lines. Cell 144: 439–452.

4. Chylicki K, Ehinger M, Svedberg H, Bergh G, Olsson I, Gullberg U. 2000. p53-mediated differentiation of the erythroleukemia cell line K562. Cell Growth Differ 11: 315–324.

5. Creyghton MP, Cheng AW, Welstead GG, Kooistra T, Carey BW, Steine EJ, Hanna J, Lodato MA, Frampton GM, Sharp PA et al. 2010. Histone H3K27ac separates active from poised enhancers and predicts developmental state. Proc Natl Acad Sci U S A 107: 21931–21936.

6. Cui K, Zhao K. 2012. Genome-wide approaches to determining nucleosome occupancy in metazoans using MNase-Seq. Methods Mol Biol 833: 413–419.

7. Davidson EH, Erwin DH. 2006. Gene regulatory networks and the evolution of animal body plans. Science 311: 796–800.

8. Ding J, Huang X, Shao N, Zhou H, Lee DF, Faiola F, Fidalgo M, Guallar D, Saunders A, Shliaha PV et al. 2015. Tex10 Coordinates Epigenetic Control of Super-Enhancer Activity in Pluripotency and Reprogramming. Cell Stem Cell 16: 653–668.

9. Dodt M, Roehr JT, Ahmed R, Dieterich C. 2012. FLEXBAR-Flexible Barcode and Adapter Processing for Next-Generation Sequencing Platforms. Biology (Basel) 1: 895–905.

10. Dong X, Weng Z. 2013. The correlation between histone modifications and gene expression. Epigenomics 5: 113–116.

11. Dreos R, Ambrosini G, Groux R, Cavin Perier R, Bucher P. 2017. The eukaryotic promoter database in its 30th year: focus on non-vertebrate organisms. Nucleic Acids Res 45: D51–D55.

12. Duncan MT, DeLuca TA, Kuo HY, Yi M, Mrksich M, Miller WM. 2016. SIRT1 is a critical regulator of K562 cell growth, survival, and differentiation. Exp Cell Res 344: 40–52.

13. Efremova M, Teichmann SA. 2020. Computational methods for single-cell omics across modalities. Nat Methods 17: 14–17.

14. Gangadharan S, Mularoni L, Fain-Thornton J, Wheelan SJ, Craig NL. 2010. DNA transposon Hermes inserts into DNA in nucleosome-free regions in vivo. Proc Natl Acad Sci U S A 107: 21966–21972.

15. Gray SM, Amezquita RA, Guan T, Kleinstein SH, Kaech SM. 2017. Polycomb Repressive Complex 2-Mediated Chromatin Repression Guides Effector CD8(+) T Cell Terminal Differentiation and Loss of Multipotency. Immunity 46: 596–608.

16. Heinz S, Benner C, Spann N, Bertolino E, Lin YC, Laslo P, Cheng JX, Murre C, Singh H, Glass CK. 2010. Simple combinations of lineage-determining transcription factors prime cis-regulatory elements required for macrophage and B cell identities. Mol Cell 38: 576–589.

17. Ito S, Shen L, Dai Q, Wu SC, Collins LB, Swenberg JA, He C, Zhang Y. 2011. Tet proteins can convert 5-methylcytosine to 5-formylcytosine and 5-carboxylcytosine. Science 333: 1300–1303.

18. Karlic R, Chung HR, Lasserre J, Vlahovicek K, Vingron M. 2010. Histone modification levels are predictive for gene expression. Proc Natl Acad Sci U S A 107: 2926–2931.

19. Klein E, Ben-Bassat H, Neumann H, Ralph P, Zeuthen J, Polliack A, Vanky F. 1976. Properties of the K562 cell line, derived from a patient with chronic myeloid leukemia. Int J Cancer 18: 421–431.

20. Kramer A, Green J, Pollard J, Jr., Tugendreich S. 2014. Causal analysis approaches in Ingenuity Pathway Analysis. Bioinformatics 30: 523–530.

21. Krausgruber T, Fortelny N, Fife-Gernedl V, Senekowitsch M, Schuster LC, Lercher A, Nemc A, Schmidl C, Rendeiro AF, Bergthaler A et al. 2020. Structural cells are key regulators of organ-specific immune responses. Nature 583: 296–302.

22. Langmead B, Salzberg SL. 2012. Fast gapped-read alignment with Bowtie 2. Nat Methods 9: 357–359.

23. Li H, Handsaker B, Wysoker A, Fennell T, Ruan J, Homer N, Marth G, Abecasis G, Durbin R, Genome Project Data Processing S. 2009. The Sequence Alignment/Map format and SAMtools. Bioinformatics 25: 2078-2079.

24. Lozzio BB, Lozzio CB, Bamberger EG, Feliu AS. 1981. A multipotential leukemia cell line (K-562) of human origin. Proc Soc Exp Biol Med 166: 546–550.

25. Lozzio CB, Lozzio BB. 1975. Human chronic myelogenous leukemia cell-line with positive Philadelphia chromosome. Blood 45: 321–334.

26. Lu S, Zong C, Fan W, Yang M, Li J, Chapman AR, Zhu P, Hu X, Xu L, Yan L et al. 2012. Probing meiotic recombination and aneuploidy of single sperm cells by whole-genome sequencing. Science 338: 1627–1630.

27. Nagel JE, Collins GD, Adler WH. 1981. Spontaneous or natural killer cytotoxicity of K562 erythroleukemic cells in normal patients. Cancer Res 41: 2284–2288.

28. Paithankar KR, Prasad KS. 1991. Precipitation of DNA by polyethylene glycol and ethanol. Nucleic Acids Res 19: 1346.

29. Prescott SL, Srinivasan R, Marchetto MC, Grishina I, Narvaiza I, Selleri L, Gage FH, Swigut T, Wysocka J. 2015. Enhancer divergence and cis-regulatory evolution in the human and chimp neural crest. Cell 163: 68–83.

30. Quinlan AR, Hall IM. 2010. BEDTools: a flexible suite of utilities for comparing genomic features. Bioinformatics 26: 841–842.

31. Rada-Iglesias A, Bajpai R, Swigut T, Brugmann SA, Flynn RA, Wysocka J. 2011. A unique chromatin signature uncovers early developmental enhancers in humans. Nature 470: 279–283.

32. Robinson JT, Thorvaldsdottir H, Winckler W, Guttman M, Lander ES, Getz G, Mesirov JP. 2011. Integrative genomics viewer. Nat Biotechnol 29: 24–26.

33. Robson MI, Ringel AR, Mundlos S. 2019. Regulatory Landscaping: How Enhancer-Promoter Communication Is Sculpted in 3D. Mol Cell 74: 1110–1122.

34. Schones DE, Cui K, Cuddapah S, Roh TY, Barski A, Wang Z, Wei G, Zhao K. 2008. Dynamic regulation of nucleosome positioning in the human genome. Cell 132: 887–898.

35. Sekhon A, Singh R, Qi Y. 2018. DeepDiff: DEEP-learning for predicting DIFFerential gene expression from histone modifications. Bioinformatics 34: i891–i900.

36. Singh R, Lanchantin J, Robins G, Qi Y. 2016. DeepChrome: deep-learning for predicting gene expression from histone modifications. Bioinformatics 32: i639–i648.

37. Smith T, Heger A, Sudbery I. 2017. UMI-tools: modeling sequencing errors in Unique Molecular Identifiers to improve quantification accuracy. Genome Res 27: 491–499.

38. Stroud H, Feng S, Morey Kinney S, Pradhan S, Jacobsen SE. 2011. 5-Hydroxymethylcytosine is associated with enhancers and gene bodies in human embryonic stem cells. Genome Biol 12: R54.

39. Stuart T, Butler A, Hoffman P, Hafemeister C, Papalexi E, Mauck WM3rd,, Hao Y, Stoeckius M, Smibert P, Satija R. 2019. Comprehensive Integration of Single-Cell Data. Cell 177: 1888-1902 e1821.

40. Stuart T, Satija R. 2019. Integrative single-cell analysis. Nat Rev Genet 20: 257–272.

41. Sutherland J, Mannoni P, Rosa F, Huyat D, Turner AR, Fellous M. 1985. Induction of the expression of HLA class I antigens on K562 by interferons and sodium butyrate. Hum Immunol 12: 65–73.

42. Teng L, He B, Wang J, Tan K. 2015. 4DGenome: a comprehensive database of chromatin interactions. Bioinformatics 31: 2560–2564.

43. Valent P, Sadovnik I, Racil Z, Herrmann H, Blatt K, Cerny-Reiterer S, Eisenwort G, Lion T, Holyoake T, Mayer J. 2014. DPPIV (CD26) as a novel stem cell marker in Ph+ chronic myeloid leukaemia. Eur J Clin Invest 44: 1239–1245.

44. Wang J, Dai X, Berry LD, Cogan JD, Liu Q, Shyr Y. 2019. HACER: an atlas of human active enhancers to interpret regulatory variants. Nucleic Acids Res 47: D106–D112.

45. Warren L, Bryder D, Weissman IL, Quake SR. 2006. Transcription factor profiling in individual hematopoietic progenitors by digital RT-PCR. Proc Natl Acad Sci U S A 103: 17807–17812.

46. Weiland G, zu Dohna HG. 1978. [The suitability of the enzyme-linked immunosorbent assay (ELISA) for the demonstration of babesia antibodies in hyperimmune serums with the use of ectoantigens]. Berl Munch Tierarztl Wochenschr 91: 33–35.

47. Welch JD, Kozareva V, Ferreira A, Vanderburg C, Martin C, Macosko EZ. 2019. Single-Cell Multi-omic Integration Compares and Contrasts Features of Brain Cell Identity. Cell 177: 1873–1887 e1817.

48. Whyte WA, Orlando DA, Hnisz D, Abraham BJ, Lin CY, Kagey MH, Rahl PB, Lee TI, Young RA. 2013. Master transcription factors and mediator establish super-enhancers at key cell identity genes. Cell 153: 307–319.

49. Williams K, Christensen J, Pedersen MT, Johansen JV, Cloos PA, Rappsilber J, Helin K. 2011. TET1 and hydroxymethylcytosine in transcription and DNA methylation fidelity. Nature 473: 343–348.

50. Yang C, Cai H, Meng X. 2016. Polyphyllin D induces apoptosis and differentiation in K562 human leukemia cells. Int Immunopharmacol 36: 17–22.

51. Yin Q, Wu M, Liu Q, Lv H, Jiang R. 2019. DeepHistone: a deep learning approach to predicting histone modifications. BMC Genomics 20: 193.

52. Young L, Sung J, Stacey G, Masters JR. 2010. Detection of Mycoplasma in cell cultures. Nat Protoc 5: 929–934.

53. Zong C, Lu S, Chapman AR, Xie XS. 2012. Genome-wide detection of single-nucleotide and copy-number variations of a single human cell. Science 338: 1622–1626.

